# Zhi-Shi-Huang-Wu slows Parkinson’s progression in transgenic *C. elegans* models

**DOI:** 10.64898/2026.03.11.709540

**Authors:** Fahim Muhammad, Yan Liu, Hui Ren, Yongtao Zhou, Hui Yang, Hongyu Li

**Author notes:** **Correspondence:** Hongyu-Li, College of Life Sciences, Lanzhou University, Lanzhou, 730000, China.

## Abstract

Parkinson’s disease (PD) is the second most progressive degenerative disorder of the brain due to dopaminergic (DA) neuron degenerations and alpha-synuclein (α-Syn) accumulations. At present, the disease has no effective treatment. Therefore, the current study objective is to identify a novel anti-PD formula (Zhi-Shi-Huang-Wu Formula, F-2) computed at 8:4:2:1 ratio from HSP 70 promoter activators Valeriana jatamansi (V), Acori talarinowii (A), Scutellaria baicalensis (S), Fructus Schisandrae (F). Traditionally, V is used to cure memory impairments, A treats mental disorders, and chronic mild stress, S for neuroprotection, and F showed multiple therapeutic actions to treat insomnia. This study investigated the neuroprotective potential of the V, A, S, F, formula F-2 and its underlying molecular mechanisms in transgenic *Caenorhabditis elegans* models. A, S, F, and F-2 successfully restored 6-hydroxydopamine intoxicated DA neuron degenerations, reduced food-sensing behavior disabilities, and attenuated α-Syn aggregations. Moreover, activates the lipid deposition and proteasome expressions to confirm α-Syn degradations at the cellular level. Reactive oxygen species (ROS) cause oxidative stress, and A, S, F, and F-2 repressed ROS and raised *SOD*-3 expressions. Overall, these data indicate that V, A, S, F combined into F-2 (22.3%) are more effective against PD progression-like symptoms than individual drugs V (0.7%), A (11.4%), S (9.6%), and F (12.6%). These improved neuroprotective actions of F-2 possibly due to following the antioxidative pathway.

**Graphical abstract:** 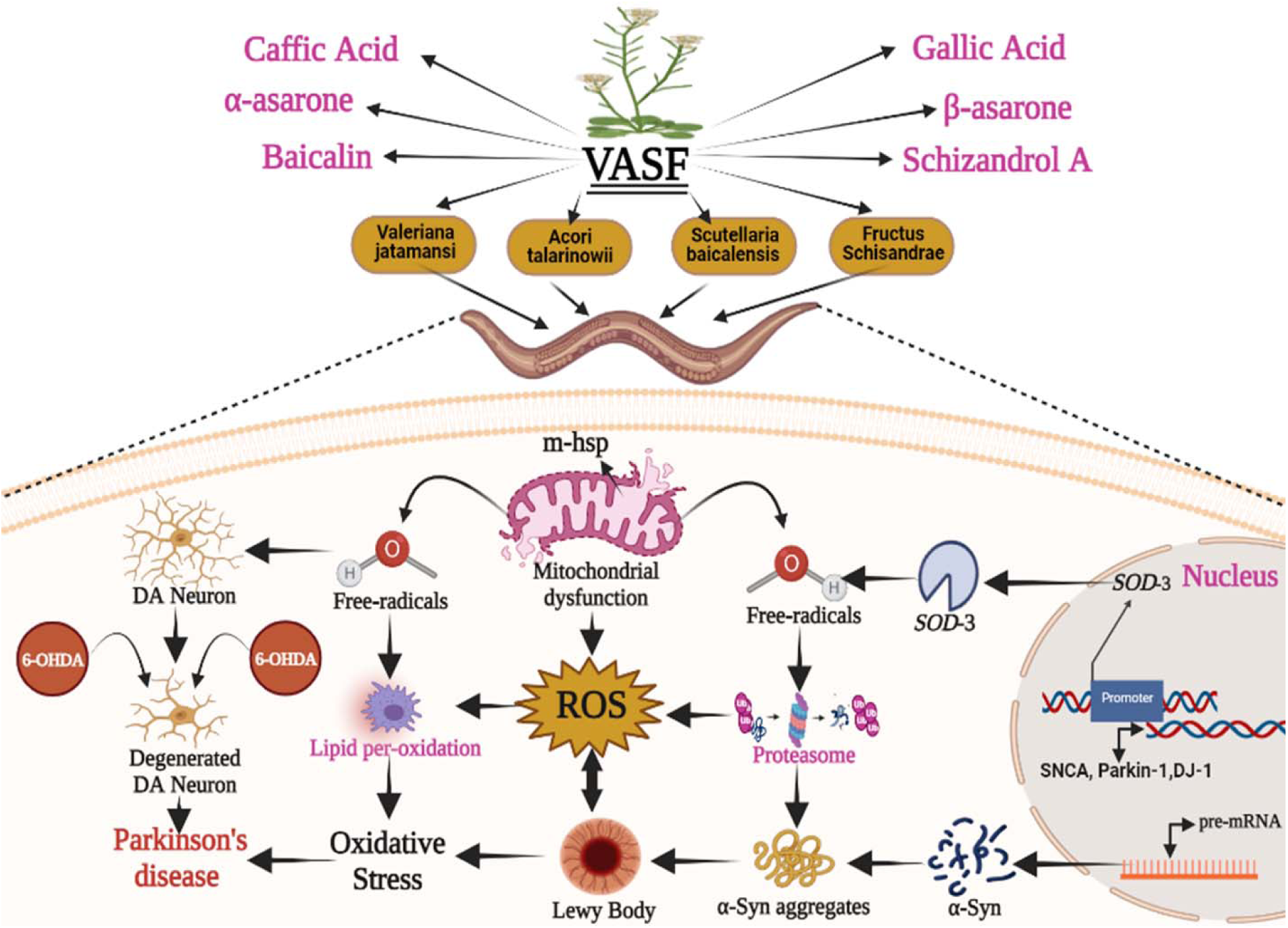

## 1 Introduction

Parkinson’s disease (PD) is the second most progressive neurological disorder after Alzheimer’s disease for decades ^1^. It is a complex degenerative disease associated with numerous motor and non-motor features. Therefore, PD patients’ motor features comprise speech difficulty, altered gait, tremors, bradykinesia, and rigidity. Whereas non-motor features are autonomic dysfunction, sensory symptoms, sleep dysfunction, and cognitive abnormalities ^2^. PD further characterized by the protein α-synuclein (α-Syn) aggregations and dopaminergic (DA) neurons degeneration in the substantia nigra pars compacta (SNPc) of the brain ^3^. Yet, PD causes remain unidentified. Current treatment for PD comprises supplementations of L-dopamine. But its long-term use causes many disadvantages in PD patients ^4^. Therefore a new therapeutic approach is to discover a plant-based neuroprotective drug to prevent and treat PD ^5^. Traditional Chinese medicine (TCM) has been cast-off for centuries. TCM has become a common and oldest medical treatment course of complementary and alternative therapeutics medicine for PD. TCM’s multi-drug, multi-target approach perfectly fits with PD’s multifactorial pathophysiology. Therapeutics derived from TCM, comprising herbal extracts and individual active components, have been used in clinics for a year’s ^6^. Therefore, we preferentially selected 35 single drugs which appeared more times by collection and statistics of the ancient Chinese prescriptions and proved to benefit neurodegenerative diseases. By screening via the ability to enhance HSP 70 promoter expressions, the top four HSP 70 promoter activator TCMs from 35 single drugs were filtered out in our laboratory. Heat shock protein can prevent false folding and protein aggregation refold the misfolded protein, especially the heat shock protein HSP 70. Studies have shown that HSP 70 plays an important role in reducing proteins accumulation of α-Syn, Aβ, restoring tau protein in vivo balance, reducing oxidative stress, and inhibiting nerve inflammation. While HSP 70 chaperone families control apoptosis at the mitochondrial stage and suppress the stress-induced signaling by inhibiting BAX translocation and enhancing the Bcl-2 expressions. HSP 70 families protected the neurons from aggregated misfolded proteins ^7^. Therefore, HSP 70 is associated with treating conformational and neurodegenerative changes such as PD. The four HSP 70 promoter activator TCM comprises Valeriana jatamansi (V), Acorus talarinowii (A), Scutellaria baicalensis (S), and Fructus Schisandrae (F). Among them, V and S is the best HSP 70 promoter activator which can activated the HSP 70 expression in pGL3-HSP70 and pRL-TK co-transfected HEK-293T cells as low as 0.4 mg/mL decoction, while others are 3.2 mg/mL (A), 1.2 mg/mL (S), 5.5 mg/mL (F), respectively.

V, with Chinese herbal name “Zhi Zhu Xiang”, its family (Caprifoliaceae) comprises more than 200 species, widespread in Europe, North America, and Asia ^8^. Many of them are well known in Chinese and Tibetan medicine as a part of various formulations to treat brain diseases ^9^. Traditionally, V is used to cure memory impairment ^8^. V is a high source of flavonoids, terpenoids, tannins, and anti-oxidants ^10^; A with Chinese herbal name “Shi Chang Pu”) is a well-known TCM which treats mental disorders. Previously, Kai Xin San’s formula, which contained A, had been proved to reduce depression-like indications of chronic mild stress ^11^. The active ingredients with antidepressant action of the A comprise of α and β-asarone (Flavonoids), and anti-oxidants are the main ingredients of A in treating brain diseases ^12^. S with Chinese herbal name “Huang Qin” is a perennial herb distributed mainly in Eastern Asia. S is the most prevalent and multi-purpose herb for neuroprotection, traditionally used in China and other oriental countries ^13^. The biological activity of the S is mainly related to flavonoid content in the roots. F with Chinese herbal name “Wu Wei Zi” is an active ingredient isolated from the fruits of Schisandra chinensis Baill. They are rich in flavonoids like Flavin glucoside and broadly used as a TCM in China, India, and Eastern Europe ^14^. F showed multiple therapeutic actions to treat insomnia, neuroinflammation, kinds of strokes, and neurodegenerations. F also exhibited various pharmacological properties to stop oxidative activities and reduce cellular apoptosis ^15^. Every drug has a long history of traditional use in different regions of China against emotional stresses, nervous disorders, insomnia, epilepsy, memory impairments, and other human diseases ^16^. Therefore, in our current study, the four HSP 70 promoter activator TCM’s (<u>supplementary</u> <u>Table:1</u>) abbreviated as V, A, S, F were selected to examine their neuroprotective effects on various PD C*. elegans* models. Transgenic *C. elegans* is an appropriate model for preliminary neurodegenerative study. The *C. elegans* have an easy culture method, a short life cycle with a simple neuron network, and a conserved nervous system pathway ^17^. Our study confirmed that drug group V enhanced worm’s lifespan, antioxidant-like properties, and lessened ROS but failed to exhibit neuroprotective effects. That’s why we have combined the drug V with A, S, F to examine whether these TCM together supported their therapeutic actions or not. We mixed V, A, F, S at an 8:4:2:1 ratio (V: 20 mg/mL, A: 10 mg/mL, S: 5 mg/mL, F: 2.5 mg/mL), respectively and computed a formula drug named “Zhi-Shi-Huang-Wu”. We found Zhi-Shi-Huang-Wu formula has significant anti-PD effects on transgenic *C. elegans* PD models. This Zhi-Shi-Huang-Wu formula was selected based on each V, A, F, S drug’s neuroprotective effect and physiological toxicity to the treated PD *C. elegans* models via orthogonal test (Supplementary Fig: S01). Later, we compared Zhi-Shi-Huang-Wu and each single drug’s V, A, F S neuroprotective functions on *C. elegans* models.

To begin with, the neuroprotective effects of every single drug and Zhi-Shi-Huang-Wu were assessed against 6-OHDA treated DA neuron and α-Syn toxicity in different PD *C. elegans* models. After that, the impact on lipid deposition, ubiquitin-like proteasome activity to degrade intracellular misfolded proteins (α-Syn), lifespan prolongation, ROS level, and the antioxidant system was tested on the V, A, S, F, and Zhi-Shi-Huang-Wu, respectively. Finally, our study would provide a new TCM formula candidate and data support for PD treatment. Our experimental assessment verified that Zhi-Shi-Huang-Wu exhibited better neuroprotective effects than single drug V, A, S, F extracts. Together, these drug in formula’s promoted and supported more effective neuroprotective actions against PD on treatment. According to our knowledge, we are the first to report this neuroprotective study of every single drug and Zhi-Shi-Huang-Wu in transgenic *C. elegans* models of PD. Besides, we will continue exploring the Zhi-Shi-Huang-Wu and further discover its other molecular pathways in future projects.

## 2 Materials and methods

### 2.1 V, A, S, F preparations

Weight every herb (V, A, S, or F) in the crude form of up to 5 grams and boil for 30 minutes twice, respectively. Separate the extract/supernatant from the residue and filter. For TCM names and originality see (**Table 1**). Then, each drug extract (supernatant) sample was prepared for freeze-drying to get drugs in powder form. After freeze-drying, quality control of crude drugs was performed by HPLC (detailed methodology is given in the supplementary data). The stock solution was prepared from the powdered drug at 50 mg/mL and diluted with distilled water to make multiple dilutions for experimental work and stored at -20 °C for later use.

**Table 1:** Selected traditional Chinese Medicine with neuroprotective properties.

| Botanical names | Scientific Names | Components | Traditional Uses | Ref |
| --- | --- | --- | --- | --- |
| Valeriana jatamansi | (nardostachyos radix et rhizoma)<br>Pharmacopoeia of China (2010, 2015) | Caffeic acid (2.80%), Valtrate (7.28%) | 1-Sedative<br>2-Anti-epilepsy<br>3-Nervous unrest | 8, 18 |
| Acori Tatarinowii | (acori tatarinowii rhizoma)<br>Pharmacopoeia of China (2015) | A-asaron (3.39%)<br>β-asarone (3.29%) | 1-resuscitate<br>2-calm the mind,<br>3-resolve dampness | 19 |
| Scutellaria baicalensis | (scutellariae radix)<br>(Japanese Pharmacopoeia, 16th edn. (2012))<br>Pharmacopoeia of China (2010, 2015) | Baicalein (2.81 %)<br>Chrysin (3.22% ) | 1-Diarrhea<br>2-Hypertension<br>3-Inflammation | 20 |
| Fructus Schisandrae Chinensis | (schisandrae chinensis fructus)<br>Hong Kong Chinese Materia Med. Standards (2010,2014) | Gallic acid (2.8 4%)<br>Schizandrol (3.19%) | 1-Liver disease<br>2-Relieve cough<br>3-Stomach Disorder | 21, 22 |

**Botanical names:** “V” represented the Valeriana jatamansi, “A” Acori Tatarinowii, “S” Scutellaria baicalensis, and “F” Fructus Schisandrae Chinensis in Table 1 above.

### 2.2 HPLC Analysis of V, A, S, F

#### 2.2.1 HPLC required Reagents

Acetonitrile 80 % (HPLC grade), and Ultrapure water (20 %) was obtained using the Milli-Q system (Millipore, Bedford, MA, USA) was used in the experiments. Methanol (HPLC grade) was used to clean the column. Eight standards (Caffeic acid, Valtrate, β-Asarone, α-Asarone, Gallic acid, Schizandrol A, Baicalin, Chrysin) for quantitative analysis were purchased from the National Institution for Food and Drug Control (NIFDC) Beijing, China.

#### 2.2.2 Analytical Conditions and Instrumentation

HPLC system 1200 series (Agilent Technologies, USA) equipped with Chemstation B.03.02 software (Agilent Technologies, USA) comprised a quaternary solvent delivery pump, an online vacuum degasser, an auto sampler, a thermostatic compartment, and a UV detector were used for chromatographic analysis. All separation processes were performed using a C_18_ column (5.0 m particle size with 250 mm ×4.6 mm i.d) Kromasil.

#### 2.2.3 Mobile phase solutions

Mobile phase A was acetonitrile, and mobile phase B was water. The linear gradient condition (30 % B for 0 min to 8 min; 30% to 20% B for 8 min to 12 min; 20% to 10% B for 12 min to 20 min; 10% B for 20 min to 30 min) was applied for the separation process. The eight selected standards can be analysed completely by using this procedure. The flow rate was 0.8 mL/min, and the column temperature was 25 ∘C, which was maintained for the entire experiment. The eluate was monitored at 314 nm, and the injection volume was 30 µL. The peak identification was based on the retention time and UV spectrum against the standard presented in the chromatogram.

#### 2.2.4 Preparation of the Standard Solution

Quantification was based on the standard external method. The stock solutions of each standard were prepared by dissolving in methanol. The solutions were separately and precisely prepared as follows: Caffic acid (0.0025 g), Valtrate (0.0025 g), β-asarone (0.0025 g), α-asarone (0.0025 g), Gallic acid (0.0025 g), Schizandrol A (0.0035 g), Baicalin (0.0025 g), and Chrysin (0.0025 g) were placed in centrifugal tubes, in which 2, 2, 2, 2, and 2.5 mL of methanol were then added, respectively. After 10 min ultra-sonication, filter with a syringe filter before analyzing the HPLC. All of the solutions were stored at −4^∘^C.

#### 2.2.5 Working standard solutions preparations

The working standard solutions were prepared using the stock solutions. The appropriate stock solutions and methanol were mixed well. Finally, the standard working solutions (800, 261.5, 255.7, 200, and 200 g/mL, 200 g/ml, 150 g/ml) of Caffic acid, Valtrate, β-asarone, α-asarone, Gallic acid, Schizandrol A, Baicalin, and Chrysin were prepared. Subsequently, the mixed standard work solutions (1.25 g/mL to 300 g/mL) of Caffic acid, Valtrate, β-asarone, α-asarone, and Gallic acid, Schizandrol A, Baicalin, and Chrysin were prepared by using the working standard solutions. All of the solutions were stored at −4 ^∘^C.

#### 2.2.6 V, A, S, F sample preparations for HPLC

The powdered samples of each drug were refluxed using ultra water for 2 hrs; this process was repeated three times at 90 °C. The extracts were combined before being filtered and then diluted with water until the final volume of 0.25 mg herbs/mL was reached. The diluted solution was filtered through a syringe filter (0.25 m) and stored at −4 °C before injection.

### 2.3 *C. elegans* culture and maintenance

Transgenic and wild-type *C. elegans* strains experimented within this research are stated in **Table 2**. All the strains were obtained from the *Caenorhabditis* Genetics Center, USA, Minnesota. All worms were kept on solid nematode growth media ^23^ incubated at 20 °C and fed with *Escherichia coli* strains OP50 as a food source, as described previously ^23^.

**Table 2:**
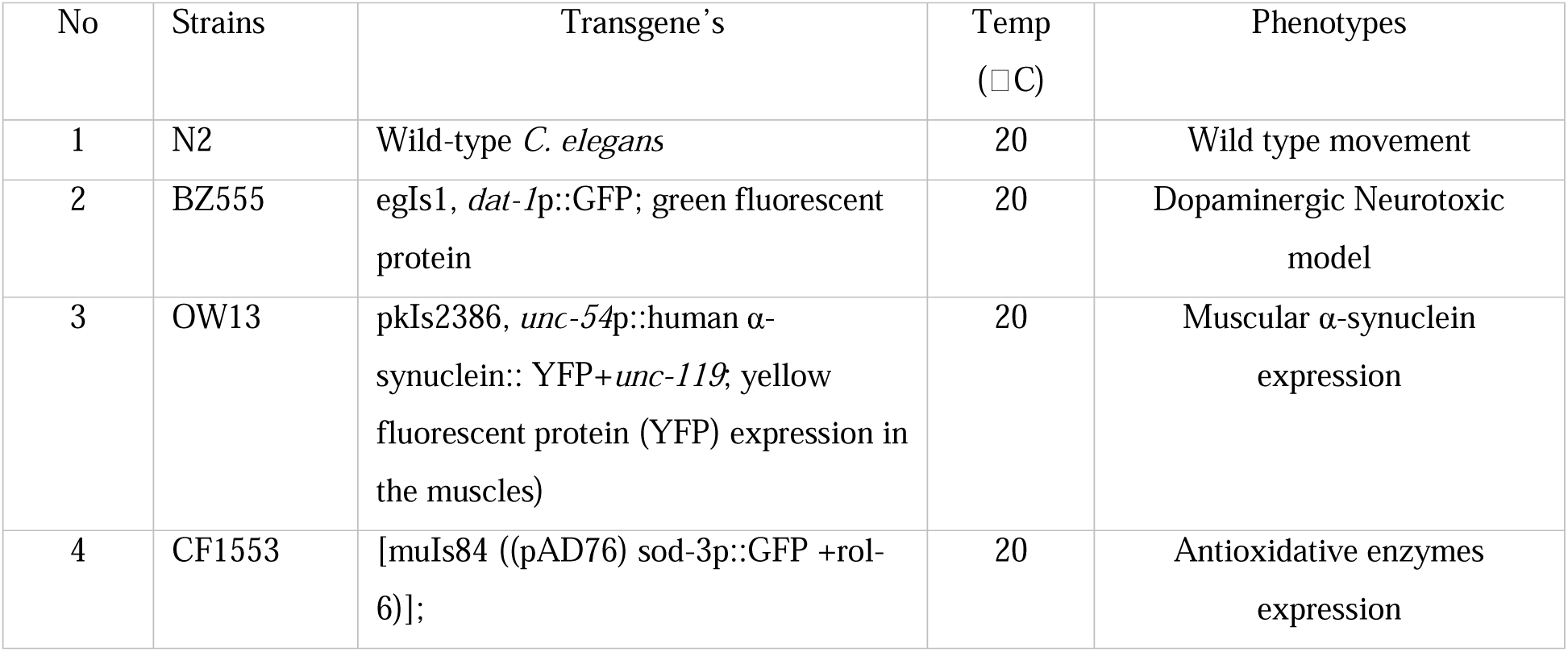
C. elegans strain used in this study and maintenance conditions.

### 2.4 Zhi-Shi-Huang-Wu formula drug Preparation

Orthogonal experiments were conducted using different combinations of V, A, S, and F according to the design shown in **Table 3**. Transgenic C. elegans strains (OW13 and CF1553) were subsequently treated with the **Table 3** listed dilutions. Following treatment and analysis, the orthogonal experimental results were used to determine the optimal formulation containing specific ratios of V, A, S, and F. The detailed results of the orthogonal experiments and the calculation procedure used to derive the formula are provided in the supplementary materials (Supplementary Figure 1). Based on these analyses, the final formulation was successfully established and named Zhi-Shi-Huang-Wu, derived from the initials of the traditional Chinese medicine components. The optimized ratio of Zhi-Shi-Huang-Wu was 8:4:2:1 (V: 20 mg/mL, A: 10 mg/mL, S: 5 mg/mL, F: 2.5 mg/mL). For convenience, this formulation is hereafter referred to as F-2.

**Table 3:** Defining the V, A, S, F drug Orthogonal experiments for evaluating the F-2 formula Percentage.

| Orthogonal experiments for F-2 formulation |  |  |  |  |
| --- | --- | --- | --- | --- |
| Groups/Cons | S (mg/mL) | F (mg/mL) | A (mg/mL) | V (mg/mL) |
| Group 1 | 0 | 0 | 0 | 0 |
| Group 2 | 0 | 2.5 | 10 | 10 |
| Group 3 | 0 | 5.0 | 20 | 20 |
| Group 4 | 2.5 | 0 | 10 | 20 |
| Group 5 | 2.5 | 2.5 | 20 | 0 |
| Group 6 | 2.5 | 5.0 | 0 | 10 |
| Group 7 | 5.0 | 0 | 20 | 10 |
| Group 8 | 5.0 | 2.5 | 0 | 20 |
| Group 9 | 5.0 | 5.0 | 10 | 0 |

### 2.5 Orthogonal Experiments for F-2 in OW13 stained with dye Nile Red and sod-3 integrated CF1553 transgenic *C. elegans*

OW13 *C. elegans* was integrated with human α-Syn protein plus YFP fusion construct, expressed α-Syn in the muscle’s walls via yellow fluorescence signals, while CF1553 integrated with *SOD*-3 gfp promoter to show superoxide dismutase (*SOD*-3) enzymatic expressions. With few modifications in the previously described protocol ^24, 25^, synchronized L1 larvae from both the worms were moved onto the NGM/OP50 plates administered with **Table 2** dilutions. Later, worms were washed with M9 buffer, mounted onto an agarose pad slide with 20 mM sodium azide, and enclosed with coverslips. Then images were observed under the fluorescence microscope (BX53; Olympus Corp., Tokyo, Japan). Later fluorescence intensity of images was measured by ImageJ software (http://imagej.net/). Assessment data is presenting in (supplementary figure 3).

### 2.6 Food clearance test

A food clearance experiment of *C. elegans* (wild-type N2, transgenic OW13, and BZ55) was performed, as previously described method ^26^. With brief modifications to conclude the appropriate, non-toxic concentration of V, A, S, F for neuroprotective treatments. Overnight grew OP50 and then suspended at a final optical density (OD) of 0.6 in nematodes S-basal solution. Every single drug was diluted at 2.5 mg/mL, 5.0 mg/mL, 10 mg/mL, 20 mg/mL, and 30 mg/mL in *E. coli* suspensions to attain the required nontoxic concentrations. Each 96-wells plate received 60 µL of the *E. coli* suspension. Randomly selected 30 L-1 worms in 10 µL of S-medium were added to an *E. coli* suspension holding a series of V, A, S, F dilutions and then incubated at 20 □C. The plate was covered with aluminium foil to prevent evaporations. The OD of the culture at 595 nm was measured once/d for 6 days using a SpectraMax M5 Microplate Reader (Molecular Devices, Silicon Valley, CA, USA). Before measuring OD, each plate was shaken for 10 sec. The fraction of worms alive per well was observed microscopically based on size. The same procedure has been done for the F-2.

### 2.7 Self-induced neurodegeneration and quantification of DA neuron via 6-OHDA treatment

BZ555 were intoxicated with 6-OHDA (Sigma, St. Louis, MI) to induce the required DA neuron degeneration. In short, with minor modifications, arranged the S-basal and added OP50 bacterial strains (S-medium). Then prepare 50 mM 6-OHDA in 10 mM ascorbic acid solution and mix with S-medium in 1.5 mL Eppendorf tube. L-3 Synchronized BZ555 worms were then moved into the experimented tube, incubated for 1 h at 20 □C, and shaken slightly every 10 min. After an hour of incubation, worms were washed three times with M9 buffer and moved to NGM/OP50 plates with or without drug V, A, S, F, or F-2 for 72 hrs. NGM/OP50 plates contained each drug at 2.5 mg/mL, 5.0 mg/mL, and 10 mg/mL and the F-2 formula. FUDR (Sigma, St. Louis, MI) was used to make *C. elegans* infertile. After inducing DA neuron degenerations and 72 hrs of drug treatment at 20 □C, BZ555 was washed three times with washing buffer (M9) and then fixed onto a 2% agar pad on a glass slide using sodium azide (20 mM) and covered with a coverslip. Immobilized BZ555 images were observed under a fluorescence microscope (BX53; Olympus Corp., Tokyo, Japan). Later the fluorescence intensities were cumulated by the software ImageJ.

### 2.8 Food-sensing behavior analysis

The DA neuron function can be identified by the movement of food sensing behaviour of *C. elegans*. Commonly, well-fed worm*s* with normal behavior exhibit a slower rate than the conditions where food is lacking. The DA neuron degenerations deteriorate the function of food sensing behavior. Therefore, first, the transgenic BZ555 and wild-type N2 *C. elegans* were treated with or without 6-OHDA for an h to induce neurodegenerations, as mentioned above. Later both the worms were treated with V, A, S, F at 2.5 mg/mL, 5.0 mg/mL, and 10 mg/mL for 72 hours. Then they were moved to new NGM plates with the presence or absence of OP50. After 5 min of calming down, the bending rate of the *C. elegans* was counted within a 20s interval. The similar procedure was used with the F-2. The slowing rate of treated and untreated *C. elegans* can be calculated using the equation provided below.

*Slowing rate = 100 – locomotory rate (%)*

*Locomotory rate (%) = Bending frequency in bacterial lawn / bending frequency outside bacterial lawn *100%*

### 2.9 Quantifications of α-syn toxicity in ow13 worms

α-Syn protein is one of the major causes of PD advancement. Therefore, we have selected transgenic OW13 worms. OW13 was integrated with human protein α-Syn plus YFP fusion construct, exhibiting α-Syn expression in the muscle’s walls via yellow fluorescence signals. With few modifications in our formerly described protocol ^27^, synchronized L-1 larvae were moved onto the NGM/OP50 plates comprising V, A, S, F at 2.5 mg/mL, 5.0 mg/mL, 10 mg/mL, and the F-2, incubated at 20 °C for 72 hrs. Later, worms were washed thrice with M9 buffer, fixed onto an agarose pad slide with 20 mM sodium azide, and covered with coverslips. Then images were observed under the fluorescence microscope (BX53; Olympus Corp., Tokyo, Japan). Later, fluorescence intensities of images were measured by ImageJ software.

### 2.10 Western blot analysis

After V, A, S, F individual drug treatment at 10 mg/mL for 72 hrs, *C. elegans* (OW13) were washed with fresh PBS and stored in a □80 refrigerator. The same procedure was applied to the F-2 formula to treat *C. elegans*. The whole *C. elegans* frozen pellets were taken the next day on the ice in lysis buffer (2X NP-40 buffer + 1 mM PMSF) and lysed with tissues grinding pestle mortar in 1.5 mL Eppendorf tubes for 40-60 sec, later heated for 10 min and centrifuge at 14000 g in 4 □C. After centrifugations, the supernatants were moved into a new tube, and the pellet was discarded. The supernatant holding the required proteins for conducting immunoblot and proteins concentration was measured by BCA kit before storing at –80 □C. Before loading the protein samples on the 16% Tricin gel for separation, protein extracts were heated for 10 minutes with a loading buffer containing SDS and DTT. After the gel run, proteins and marker bands were transferred to PVDF membranes. Then blocked the protein bands with 10% skimmed milk in 1X TBST for two hrs. After careful washing, the primary antibody (human α-Syn, mouse monoclonal, 1:3,000; LB509, Zymed, San Francisco, USA) was used for the whole night at 4 □C. The next day washed the membrane with 1X TBST for 5 minutes thrice and then used a secondary antibody (goat-anti mouse, 1:5,000; BioRad, Hercules, CA, USA) for 1:30 hrs at room temperature. Binding was visualized by the horse-radish peroxidase chemiluminescence (ECL Lumi-Light, Roche, Germany). Bands images were quantified by ImageJ software.

### 2.11 Quantification of lipid deposition

In OW13, lipid depositions were stained by Nile red dye. With minor modifications in the formerly described protocol (Jadiya et al., 2011). The stock solution was prepared by liquefying 0.5 mg Nile red dye in 1 mL acetone, then mixed at a ratio of 1:250 with *E. coli* OP50 suspensions. Later, OW13 at the L-1 were cultured on NGM/ Nile red/OP50 plates with V, A, S, F at 2.5 mg/mL, 5.0 mg/mL, and 10 mg/mL separately and in a combined F-2 group at 20 °C for 72 hrs. The control groups were treated without drugs. Finally, OW13 were washed thrice with M9, mounted onto agar pads using 20 mM sodium azide, and mounted with a coverslip. The OW13 imaged were observed in a fluorescence microscope to identify the lipid deposits. The fluorescent intensities of images were measured via ImageJ software.

### 2.12 Ubiquitinated-like proteasome activity in *C. elegans*

The activity of ubiquitin-like proteasomes in OW13 was examined using a formerly defined protocol ^28^. Worms were treated with V, A, S, F drug groups at 10 mg/mL one by one and F-2. Control groups were treated without drugs. With slight modifications, worms were lysed on ice in proteasomes lysis solution NP-40 and 1mM PMSF via tissues grinding pestle mortar for homogenizations. Later, lysates centrifugation was performed at 14,000 g at 4 □C for 10 min, then transferred the supernatants into a separate 1.5 mL Eppendorf tube containing the proteasome subunits plus protein and discarded the pellets. From supernatants containing tube, 20 μL was loaded on a 96-wells plate consisting of fluorogenic substrate up to 30 μL for each test. To observe the expressions of ubiquitin-like proteasomes in the worm’s lysate, we used LLVY-R110-AMC (Sigma-Aldrich, MAK172-IKT, USA) as the fluorogenic substrate. Proteasome cleavages the LLVY-R110 substrate and generates a green-fluorescent signal on R110 separations. The lysate and fluorogenic substrate were incubated for 1.5 hrs at 37 □C in black 96 wells microtiter plates. Then, reading was recorded at 490 nm excitation wavelengths and 525 nm emissions wavelengths at 10 min intervals for 1 hour by M2 Microplate Reader SpectraMax (Molecular Devices, Silicon Valley, CA, USA).

### 2.13 Reactive oxygen species assay

Two complementary approaches were used to evaluate the antioxidant effects of V, A, S, F, and the combined formula F-2 in wild-type N2 and transgenic BZ555 *C. elegans*. The assay was adapted from a previously reported protocol with minor modifications ^29^. For the first method, intracellular ROS in N2 worms was measured using the H_2_DCF-DA fluorescence assay. Briefly, N2 worms were first exposed to 6-OHDA to induce neurodegeneration similar to that observed in toxin-treated BZ555 worms. The 6-OHDA–treated animals were then incubated with V, A, S, or F (10 mg/mL) or the F-2 formula at 20 °C for 72 h. After treatment, live worms were washed three times with M9 buffer, and at least 40 worms were transferred to each well of a 96-well plate containing 200 µL PBS with 50 µM H_2_DCF-DA. Fluorescence was recorded every 20 min for 200 min (excitation 485 nm, emission 520 nm) using a SpectraMax M2 microplate reader (Molecular Devices, CA, USA).

In the second approach, intracellular ROS levels were quantified in BZ555 worm lysates. Following treatment with 6-OHDA and the respective compounds, worms were washed with PBS containing 1 mM PMSF and stored at −80 °C overnight. Worm pellets (∼0.04 g, approximately 1000 worms) were thawed and homogenized in 50 µL lysis buffer (PMSF and NP-40) using a motorized pestle on ice. The lysates were centrifuged at 14,000 g for 10 min at 4 °C, and the supernatants were collected. A 20 µL aliquot of the supernatant was mixed with 30 µL H_2_DCF-DA in a black 96-well plate to achieve a final concentration of 50 µM. Plates were incubated in the dark for at least 1 h, and fluorescence was measured every 10 min for 100 min (excitation 490 nm, emission 525 nm) using a SpectraMax M2 microplate reader (). The F-2 formula was evaluated using the same procedure.

### 2.14 Antioxidant superoxide dismutase (*sod*-3) assay

The antioxidant assay was performed with minor changes in the previously described method ^30^. We used transgenic strains CF1553, which strongly expressed *SOD*-3::GFP. To begin with, we treated synchronized L-1 CF1553 with V, A, S, F, at 10 mg/mL and F-2 for 72 hrs and incubated at 20 □C. After 72 hrs incubation, CF1553 were washed with M9 buffer, mounted on an agarose pad with 20 mM sodium azide in 10 µL, and enclosed with a coverslip. The V, A, S, F, and F-2 treated or untreated CF1553 worm’s fluorescence intensity was measured by Olympus, BX53, Japan fluorescence microscope. Later images intensities were calculated via ImageJ software. For positive control groups, CF1553 were treated with Juglone at 50 µM and incubated at 25 □C for 30 min.

### 2.15 Lifespan analysis

PD is an age-related degenerative disorder. Therefore, we have performed a lifespan test with a simple modification in the formerly described protocol to examine the V, A, S, F, and F-2 effect on worm age. Both BZ555 and wild-type N2 *C. elegans* were exposed to 50 mM 6-OHDA to induce self-neurodegenerations like the BZ555 neurotoxin treatment, as mentioned earlier. Then, the 6-OHDA-exposed *C. elegans* were treated with or without V, A, S, F drugs at 10 mg/mL for 72 hrs at 20 □C. Later, it washed with M9 and moved into new NGM plates seeded with strains OP50 and incubated at 20 □C until all *C. elegans* became dead. Each *C. elegans* was marked as live, dead, or censor at day intervals, considered by their pharyngeal propelling. The *C. elegans* that execute normal behaviour or have their pharynges pumping were considered alive. The *C. elegans* was scored dead because the pharyngeal pump stopped working by gentle touch. Besides, the worm displaying bizarre behaviours, with unusual death, or skulking out of the plate, was considered a censor and omitted from the calculations. On the other hand, the observations were examined until all the *C. elegans* became dead and the lifespans compared to the control groups.

### 2.16 Statistical analysis

Statistical analysis was done using Graph Pad PRISM, Version 8.02 (1992–2019 Graph Pad Software, Inc., La Jolla, CA, and the USA). In all the treated GFP tagged worms the fluorescence intensities were observed under a fluorescence microscope (BX53; Olympus Corp., Tokyo, Japan) and quantified by using ImageJ software. Each experiment was repeated thrice and normalized with control groups. Examination of significance differences (*p*-values=*p* < .001 to p< .005) by SPSS (one-way) ANOVA, followed by Tukey’s or Dunnett’s (where comparison was only to the untreated groups)ur post hoc tests were applied.For all experiments, *p*≤.005 is considered statistically significant.

## 3 Results

### 3.1 V, A, S, F, and F-2 HPLC

For detailed results, please refer to the supplementary data file (supplementary figure 1<u>, and</u> supplementary figure 2)

### 3.2 F-2 of V, A, S, F is the best combination formula on α-Syn accumulated toxicity, enhancing lipid depositions and *SOD-3* expressions

Based on the food clearance experiment results, the highest non-toxic concentration of V, and A were at 20 mg/mL, and S, F was at 10.0 mg/mL. Thus, the Orthogonal experiment was performed according to the drug dilutions in **Table 2**. To find out the best combination of V, A, S, F for PD treatment, a series combination of V, A, S, F (G-1 G-9) to treat OW13 *C. elegans*. From the results given in the (supplementary figure 3A and 3B), we concluded that the V, A, S, F from G-2 G-9 dilutions significantly reduced the α-Syn aggregated toxicity G-2 (34.3%), G-3 (40.6%), G-4 (36.1%), G-5 (29.4%), G-6 (33.5%), G-7 (36.2%), G-8 (67.3%), G-9 (55.4%).

Later we have treated G-1 G-9 on OW13 stained with dye Nile Red and CF1553 transgenic worms integrated with sod-3 genes in a similar way (Supplementary Figure 3). On conclusion’s we found V, A, S, F dilutions at 8:4:2:1 ratio’s representing V: 20 mg/mL, A: 10 mg/mL, S: 5 mg/mL, F: 2.5 mg/mL that verified to be nontoxic and highly effective in reducing α-Syn accumulations and enhancing antioxidants named as formula (Zhi-Shi-Huang-Wu, F-2) in this study. The further orthogonal experimental study of lipid depositions and *SOD*-3 expression is mentioned in the supplementary data file (Supplementary Figure 3).

### 3.3 Food Clearance Test determined the Suitable, non-toxic doses of V, A, S, F and F-2 on N2, OW13, and BZ555 *C elegans*

To determine the effective and nontoxic drug concentration of V, A, S, F extract, we performed a food clearance assay in *C. elegans* models of PD. OW13, BZ555 and N2 *C. elegans* were treated with V, A, S, F at 0 mg/mL, 2.5 mg/mL, 5.0 mg/mL, 10, 20 mg/mL, and 30 mg/mL concentrations in S-medium. *C. elegans* treated with F and S at 20, and 30 mg/mL dilutions indicated toxic behaviour, causing the abridged body sizes with fewer offsprings, even causing the death of adults at day’s intervals and dropping the food clearance curve. The remaining drug extract-treated *C. elegans* groups were active up to 10 mg/mL **(Figure 1)**. Therefore, the *C. elegans* were treated with nontoxic V, A, S, F dilutions up to 10 mg/mL. Similarly, the non-toxic concentration results of the F-2 was measured in the similar way as described above. F-2 results confirmed no toxicity effect on growth and feeding behavior of treated worms.

**Figure 1:**
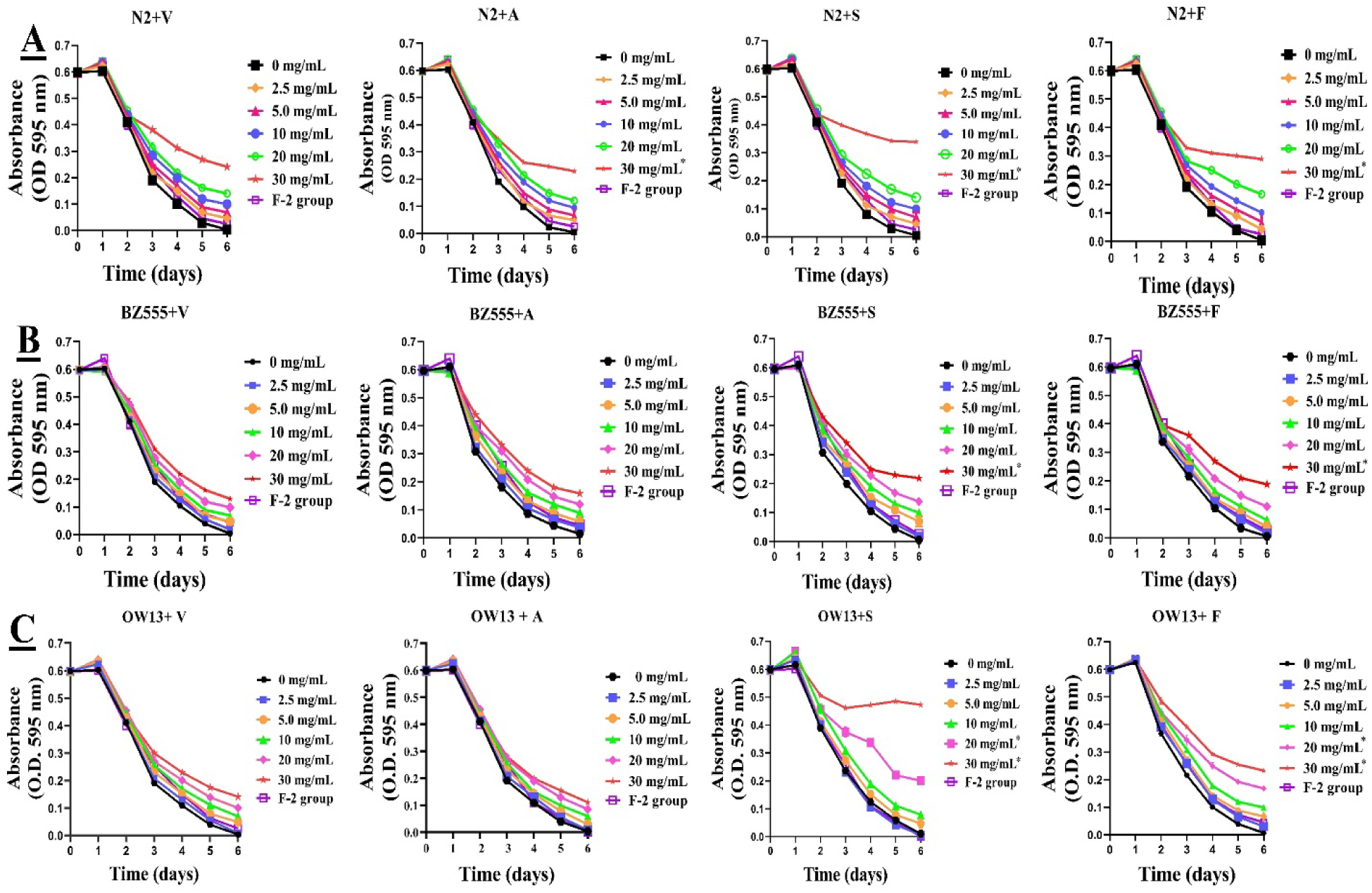
Explains the food clearance test results of V, A, S, F, and F-2 extract in the pharmacological *C. elegans* models. A food clearance test is performed to evaluate the non-toxic concentrations of selected drugs against treated models. Briefly, in a 96-well plate, synchronized L1 worms of wild-type N2, transgenic BZ555, and OW13 *C. elegans* were cultured with OP50 bacterial strain (OD A_595_ = 0.6 absorbance’s) feeding medium containing drugs V, A, S, F at 0, 2.5 mg/mL, 5.0 mg/mL, 10 mg/mL, 20 mg/mL, and 30 mg/mL concentrations and F-2 group for 6 days. The OD value of each treatment was measured and recorded on the day interval (n=10). Results showed that only F and S at 20 mg/mL and 30 mg/mL showed drug toxicity and caused *C. elegans* to become thin and slender in growth, even leading to death. All the selected V, A, S, F drugs showed non-toxic dilutions up to 10 mg/mL and the F-2 drug group to treat *C. elegans* Fig 1**. (A, B, & C)**. While the significant difference between the V, A, S, F dilutions in treated models (N2, BZ555, OW13) was not found except for drug group V and F at 20 mg/mL and 30 mg/mL *p*≤0.005 (**).

### 3.4 F-2 diminished 6-OHDA-treated da neuron degenerations better than V, A, S, F

BZ555 strains are used as a PD model integrated with DA neurons tagged dat-1 promoter (Pdat-1: GFP), which encodes for the DA transporter. DA neurons of the transgenic strain were observed under the fluorescence microscope, particularly the two pairs of cephalic sensilla’s (CEPs) at the head regions, illuminated by green colour. Selective degenerations of DA neurons by 6-OHDA showed lower fluorescence intensity when observed under the microscope. In 6-OHDA exposed *C. elegans*, the fluorescence expression decreased approximately by 67.667 % (*p*≤ 0.005) compared to the control groups, approving the neurotoxin’s degenerative effects (**Figure 2A)**. On V, A, S, F treatment at 2.5, 5.0, and 10.0 mg/mL, the fluorescence intensity of 6-OHDA intoxicated DA neurons in BZ555 raised compared to the untreated group (**Figure 2B)**. The V group is the only drug group that did not show satisfactory results on treatment as compared to the A, S, F groups (**Figure 2A&2B)**. While A, S, F diminished the 6-hydroxydopamine (6-OHDA) intoxicated DA neurons degeneration and recovered damaged neurons up to 45.57% (A), 63.62% (S), and 59.9% (F) compared to control groups and 22.41% (A), 21.61% (S), 20.71% (F) than BP in BZ555 worms. BP here used as a positive control group.. Further, we have treated 6-OHDA exposed *C. elegans* with the F-2 to verify the effects of the combined drugs in the form of formula against degenerated DA neurons recovery after 6-OHDA treatment. Our formula F-2 significantly recovered the 6-OHDA degenerated DA neurons by about 78.5 % (*p*≤ 0.005) compared to the control groups **(Figure 2B).** These results showed V, A, S, F in F-2 supported and enhanced their neuroprotective actions against PD.

**Figure 2:**
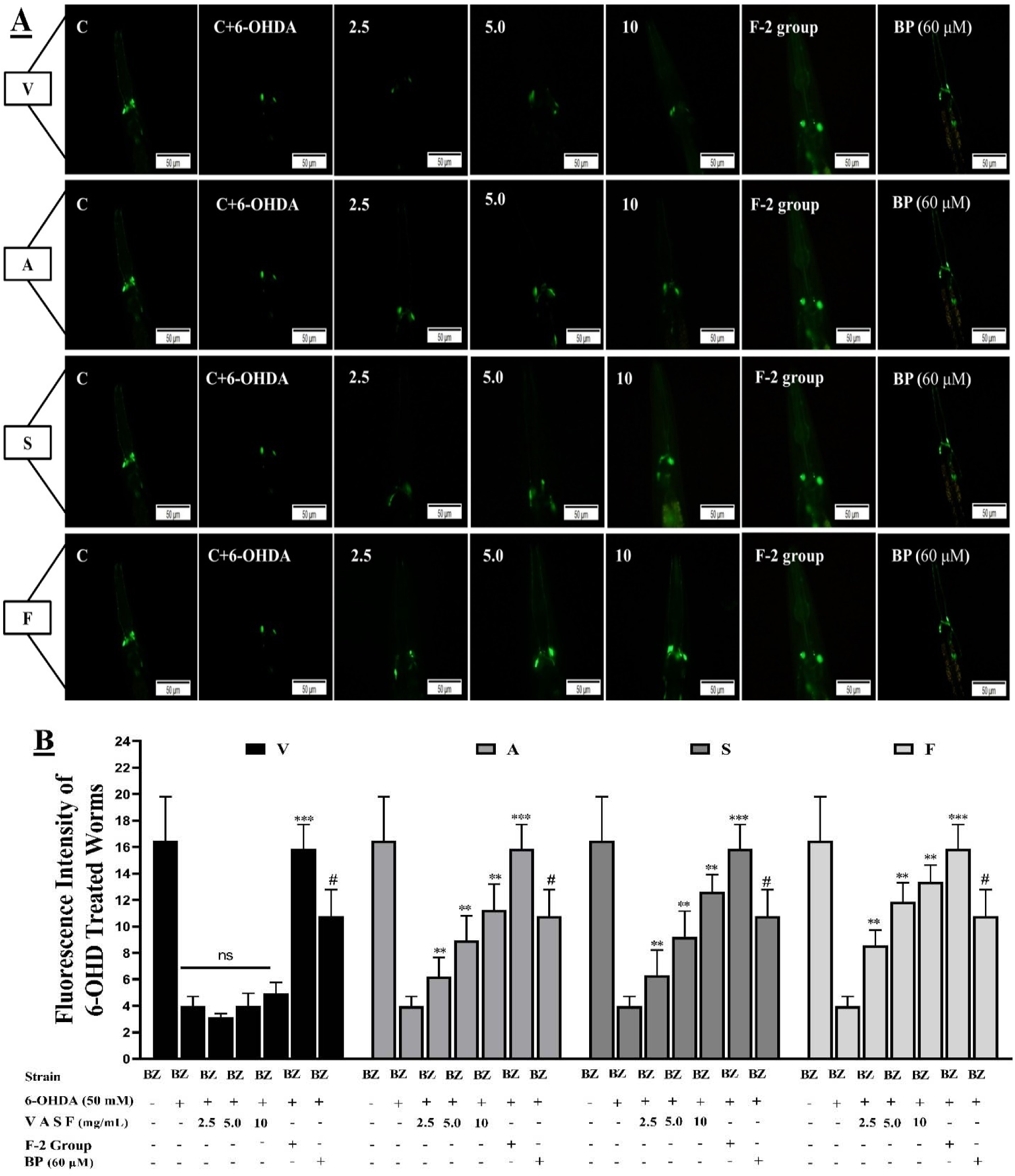
(A&B) F-2 enhanced the recovery of self-neurodegenerative DA neurons via 6-OHDA better than V, A, S, F in BZ555. (A) Images showing thefluorescence intensities of BZ555 worms treated with 6-OHDA/ (V,A, S, F) or without 6-OHDA/ (V, A, S, F) at 2.5 mg/mL, 5.0 mg/mL, 10 mg/mL and F-2. The scale bar was 50 μm. (B) The graphical demonstrations of V, A, S, F and F-2 treated drugs in 6-OHDA degenerative DA neurons. Three drug groups A (45.57%), S (63.62%), and F (59.9%) compared to control groups and A (22.41%), S (21.61%), F (20.71%) than BP in BZ555 worms showed effective results in recovering damaged DA neurons. Drug group V did not augment the fluorescence intensity in BZ555 on treatment. Besides, treatment with the F-2 (78.5 %) showed more effective results against 6-OHDA degenerative DA neurons than V, A, S, F. Data calculated by mean±SD (n = 3). ns showing that there is no significant difference between V drug-treated groups and control groups. While ** indicated the significant difference of p≤0.005 between the 6-OHDA/Cont groups and 6-OHDA/A, S, F treated groups, while *** represented the significant difference of p≤0.005 between the 6-OHDA/F-2 and the 6-OHDA/Cont groups. # represented the significant difference between n-Butylidenephthalide (BP, 60 μM) treated groups with 6-OHDA/Cont groups (p≤0.005). BP here is used as a positive control group.

### 3.5 F-2 restored food-sensing behavior in 6-OHDA-treated *C. elegans* greater than V, A, S, F

The *C. elegans* can sense foods sources while skulking on the NGM media plates. The interceded DA neuron circuits control the *C. elegans* slowing rate via sensing the foods. 6-OHDA degenerative DA neurons decrease the dopamine levels, causing the *C. elegans* to be defective in perceiving foods. We observe whether V, A, S, F and F-2 could restore the food-sensing behavior of *C. elegans* after 6-OHDA exposure or not. Our experimental assessment showed that the bending frequency of N2 generally declines in the bacterial lawns compared to 6-OHDA intoxicated *C. elegans*. Next, we examined 6-OHDA exposed N2 and BZ555 *C. elegans* to observe liabilities in food-sensing (counted as the "slowing rate"). The average slowing rate of N2 and BZ555 *C. elegans* was 89.473%. Similarly, it decreases expressively in both the worms (N2 & BZ555) treated with 6-OHDA, indicating a DA neurons degeneration causes a deficiency in food-sensing behaviour (**Figure 3A&3B)**. When treated with V, A, S, F at 2.5 mg/mL, 5.0 mg/mL, 10 mg/mL, the slowing rate increases significantly compared with 6-OHDA-intoxicated *C. elegans* (N2, and BZ555) up to 74.23%, *p*≤0.005 (A), 67.411%, *p*≤0.005 (S), 76.46%, *p*≤0.005 (F), and 73.551%, *p*≤0.005 (A), 71.33%, *p*≤0.005 (S), 70.423%, *p*≤0.005 (F) respectively, in dose-dependent manners. The V group didn’t show effective results in recovering degenerated DA neurons compared to the other three drug groups (A, S, and F). Formula F-2 confirmed that V, A, S, F in combination could restore 6-OHDA mediated DA neurons dysregulations much better than individual drugs. In conclusion F-2 (84.457%, *p*≤0.005) synergises their neuroprotective actions in recovering degenerative DA neurons.

**Figure 3:**
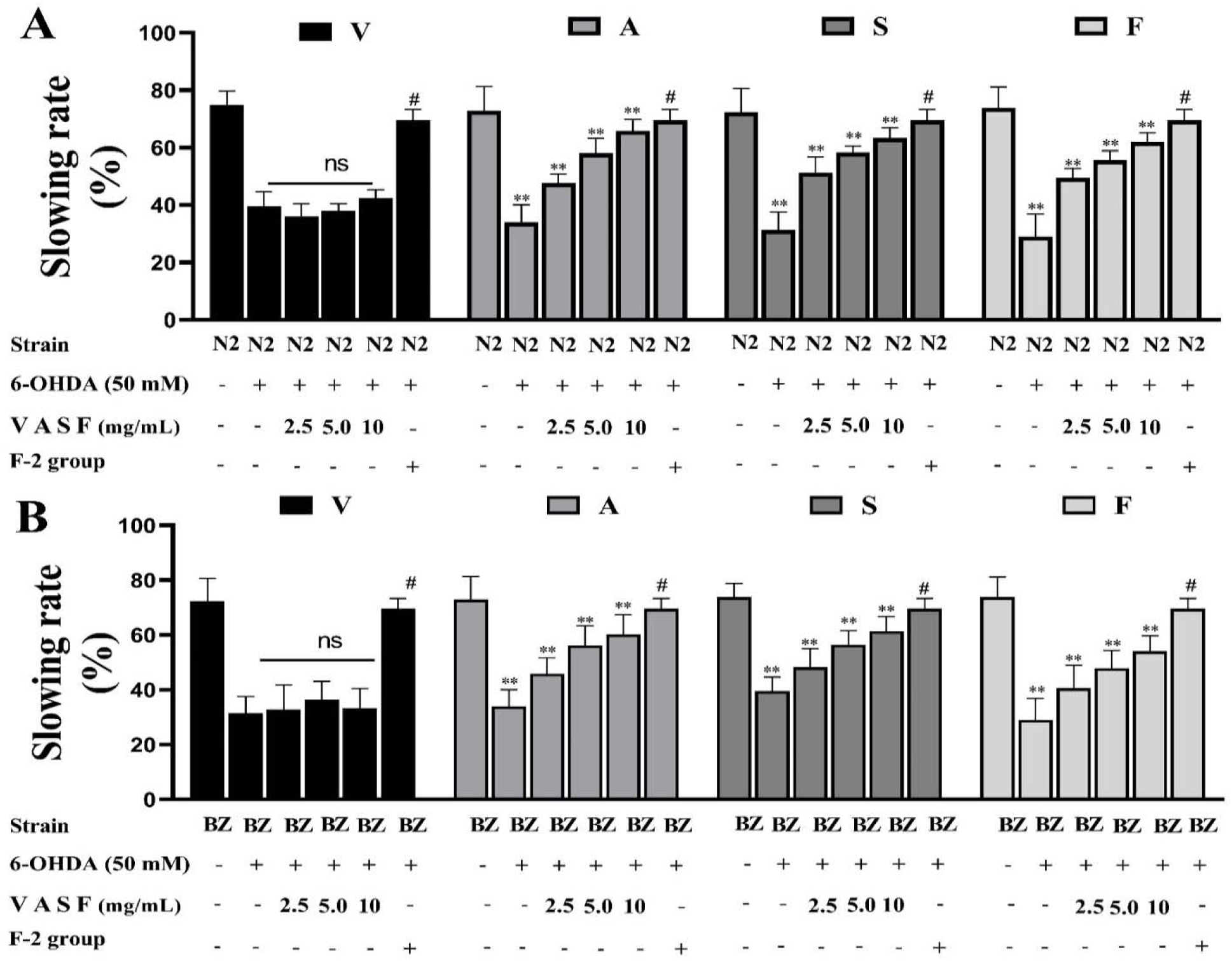
(A&B) Depicting the slowing rates of wild-type N2 and transgenic BZ555 C. elegans. The locomotory rate of both the *C. elegans* treated with 6-OHDA/V, A, S, F or without 6-OHDA/V, A, S, F at 2.5 mg/mL, 5.0 mg/mL, 10 mg/mL and F-2 for 72 hrs in the bacterial lawn or outside the bacterial lawn was evaluated in this food sensing test. Drug group V did not improve the worm’s behavior than the other three (A, S, F). At the same time, F-2 showed effective results in degenerative DA neuron recovery and improved the worm’s behavior toward perceiving foods (OP50). Data calculated by mean±SD and percentage of increase (n = 3). ns showing that there is no significant difference between V drug-treated groups and control groups. ** indicated the significant difference of 74.23%, *p*≤0.005 (A), 67.411%, *p*≤0.005 (S), 76.46%, *p*≤0.005 (F) *p*≤0.005 in N2 depicting via (A) and BZ555 treated with 73.551%, p≤0.005 (A), 71.33%, p≤0.005 (S), 70.423%, p≤0.005 (F) at 2.5 mg/mL, 5.0 mg/mL, 10 mg/mL groups vs 6-OHDA/Cont groups (B). At the same time, # indicated the significant difference between 6-OHDA/F-2 (84.457%, p≤0.005) treated and 6-OHDA/Cont group’s, *p*≤0.005.

### 3.6 F-2 attenuate the α-Syn aggregations greater than V, A, S, F in transgenic *C. elegans*

To examine the expressions and aggregations of α-Syn, we employ the transgenic strain OW13, which was created with human protein α-Syn fused with a green fluorescent protein (GFP) under the control of the unc-54 promoters where genes expressions arose in the body walls of muscle cells ^31^. OW13 strain’s advantages lead to the high efficiency of expressions and the PD-like progressive defects of motility in *C. elegans*. Therefore, it demonstrated the in vivo aggregation of α-Syn toxicity ^32^. Initially, we treated the worms with V, A, S, F individually to analyze the effects of drugs via unc-54p: YFP expressions. The data were presented as the average intensity per body area. We found that A, S, F at 2.5 mg/mL, 5.0 mg/mL, 10 mg/mL in OW13 worms showed significant effects in reducing the α-Syn accumulated toxicity than group V compared to untreated groups. The fluorescent intensities of YFP expression in OW13, representing α-Syn protein expressions, significantly decreased by about 26% (A, *p*≤0.005), 41 % (S, *p*≤0.005), and 34% (F, *p*≤0.005), and 14.66 % (p≤ 0.005) than n-Butylidenephthalide (BP) compared with untreated groups. PB was used as a positive control group. Next, we treated OW13 worms with the F-2. F-2 showed superior results in decreasing the aggregative toxicity of α-Syn up to 49.88 % (*p*≤0.0005) than un-treated groups. While 34.6 % greater attenuated effect than BP. Hence, we can say that F-2 supported and enhanced their protective actions against α-Syn accumulated toxicity more effectively than the single drug group (**Figure 4A& 4B)**.

**Figure 4:**
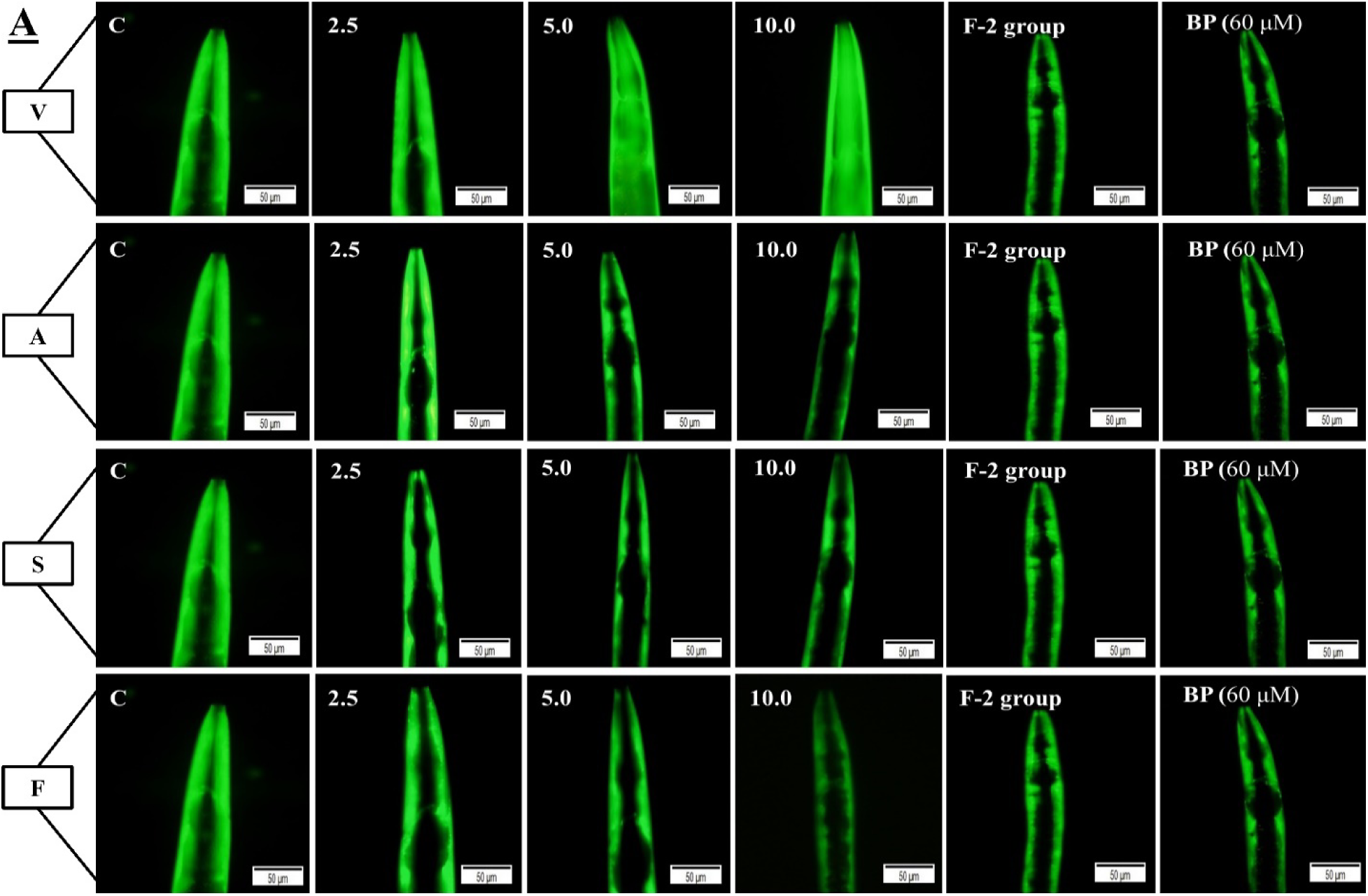
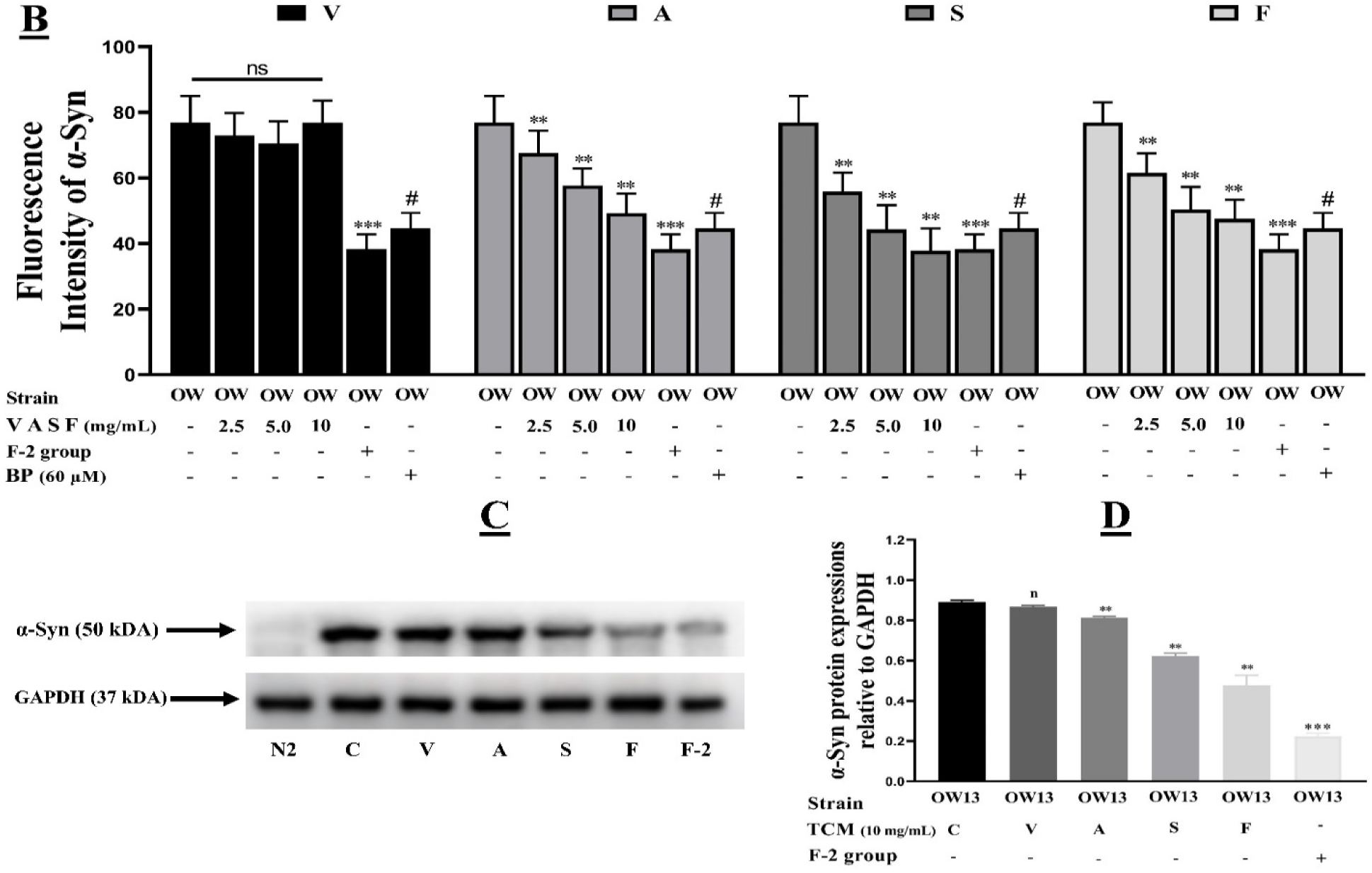
**(A&B)** Drug group F-2 on treatment reduced the α-Syn accumulated toxicity much better than V, A, S, F, in transgenic worms OW13. **(A)** GFP fluorescence expressions in the muscles of worms OW13. The scale bar was 50 μm. **(B)** Bar Graph representing the quantitative data of the fluorescence intensities of V, A, S, F and F-2 in OW13. Drug group V did not reduce the α-Syn accumulation in the worm’s muscles on treatment. ImageJ software was used to enumerate the fluorescence intensities of images. Data were computed by mean±SD (n = 3). ns depicting no significant difference between drug V dilutions with control groups. Similarly, ** shows a significant difference between the drugs A (26%), S (41 %), and F (34%) treated, and untreated groups (*p*≤0.005), and *** indicated a significant difference of *p*≤0.005 between the F-2 (49.88 %) and the untreated groups. While (#) showing a considerable difference of *p*≤0.005 between the positive control group BP (60 μM) with the untreated (control) groups. **(C)** V, A, S, F and F-2 reduced the α-Syn protein expression by the immunoblot assessment. Results showed that A, S, F significantly reduced the α-Syn accumulations in OW13 worms than group V. In combination, the F-2 (61.54%, *p*≤0.005) reduced α-Syn aggregations much higher than single drug’s A (13%, *p*≤0.005), S (24%, *p*≤0.005), F (41%, *p*≤0.005). **(D)** Graphical effects of immunoblot band intensities of V, A, S, F in single and F-2 relative to the GAPDH (37 kDa). ns shows no significant difference between V drug-treated groups relevant to GAPDH. ** depicting the significant difference of (*p*≤0.005) between A, S, F treated and untreated groups. # indicated the significant difference of (*p*≤0.005) between F-2 drug and untreated groups.

### 3.7 F-2 reduced α-Syn accumulation greater than V, A, S, F confirmed via western blotting

Western blotting detected the α-Syn protein accumulations ^33^. To further verify the V, A, S, F and F-2 effects in decreasing proteins α-Syn aggregative toxicity in OW13. We have treated OW13 worms with V, A, S, F at 10 mg/mL for 72 hrs, and western blotting was assessed using antibodies (primary and secondary). Besides, we treated OW13 worms with F-2 implemented the same protocol. GAPDH (37 kDa) was used as a positive control, and data were measured with GAPDH protein expressions relative to drug-treated and untreated groups. Western blot results proved that drugs group A (13%, *p*≤0.005), S (24%, *p*≤0.005), F (41%, *p*≤0.005) on treatment reduced α-Syn accumulated expressions relative to the GAPDH groups. The V on treatment did not show significant reductions of protein α-Syn accumulations relative to the GAPDH. In contrast, F-2 showed significantly better results in lowering α-Syn accumulation relative to the GAPDH by about 61.54%, *p*≤0.005 (**Figure 5A&5B)**.

**Figure 5:**
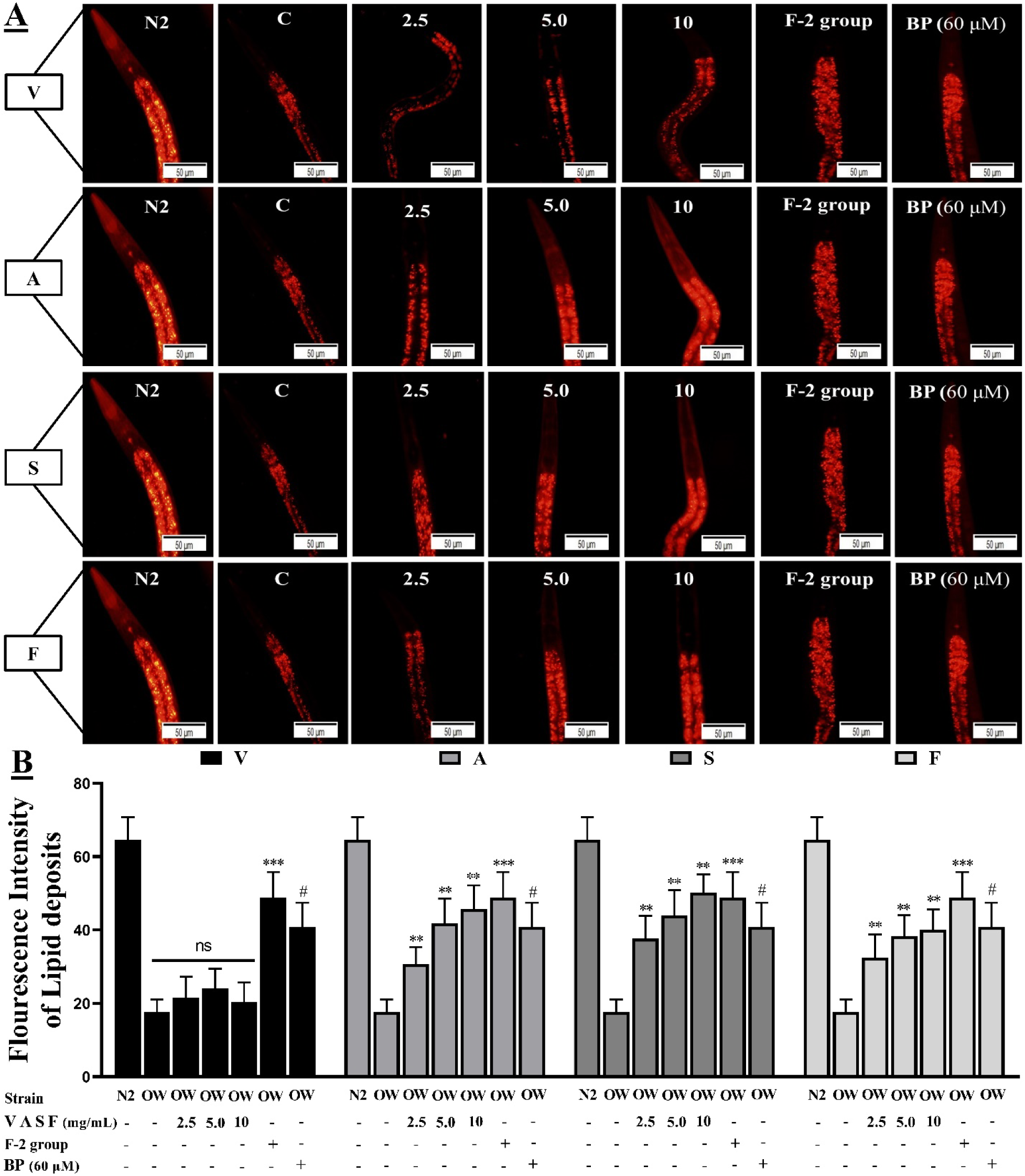
**(A&B)** F-2 increased the lipid deposits higher than V, A, S, F drug groups in OW13. **(A)** Representative images are the Nile red staining of OW13 worms after V, A, S, F at (2.5, 5.0, 10.0) mg/mL treatment for 72 hrs. The scale bar was 50 μm. **(B)** A graphical picture of fluorescence intensities of OW13 was stained with the Nile red and quantified by software ImageJ. The data was calculated by mean ± SD (n = 3). ns depicting no significant difference compared with control and V drug dilutions. Similarly, drugs group A (43%, *p*≤0.005), S (38%, *p*≤0.005), F (44%, *p*≤0.005) showed a significant difference of *p*≤0.005 compared to the control groups represented by **. While *** describes the significant difference of *p*≤0.005 between the F-2 with untreated groups. Similarly, # means the significant difference of *p*≤0.005 with **BP** (60 μM) and the untreated groups.

### 3.8 F-2 rrecovered lipid deposition much higher than V, A, S, F in OW13 *C. elegans*

α-Syn has associations with fatty acid modifications and lipids molecule in which it starts vesicle formations through specific mechanisms ^34^. In OW13, the lipids content significantly decreased because of α-Syn accumulations and interrupted its ubiquitin-like lipids depositions. Moreover, the toxic nature of aggregated α-Syn tends to augment lipids peroxidation. It produces ROS, which is another major cause of PD. We treated the OW13 with V, A, S, F at 2.5 mg/mL, 5.0 mg/mL, and 10 mg/mL and then used Nile red staining method to detect the lipid deposits. We have conducted experiments to observe the lipid level in OW13 worms under both V, A, S, F treatment and non-treatment. Drug groups A (43%, *p*≤0.005), S (38%, *p*≤0.005), F (44%, *p*≤0.005) significantly enhanced the lipid level dose-dependently on treatment except for the drug groups V (**Figure 6A&6B)**. Next, we have treated the transgenic *C. elegans* with F-2. Our results showed F-2 (57%, *p*≤0.005) improved their anti-α-Syn actions and increased the lipid content in OW13 worms than control groups and similarly by about 24.331% more than the average of individual drug groups (A, S, F) on treatment. Further we have treated the worms with BP, and our results showed BP augmented the lipid contents up to 34.67 % than untreated group. BP (60 µM) was used here as a positive control.

**Figure 6:**
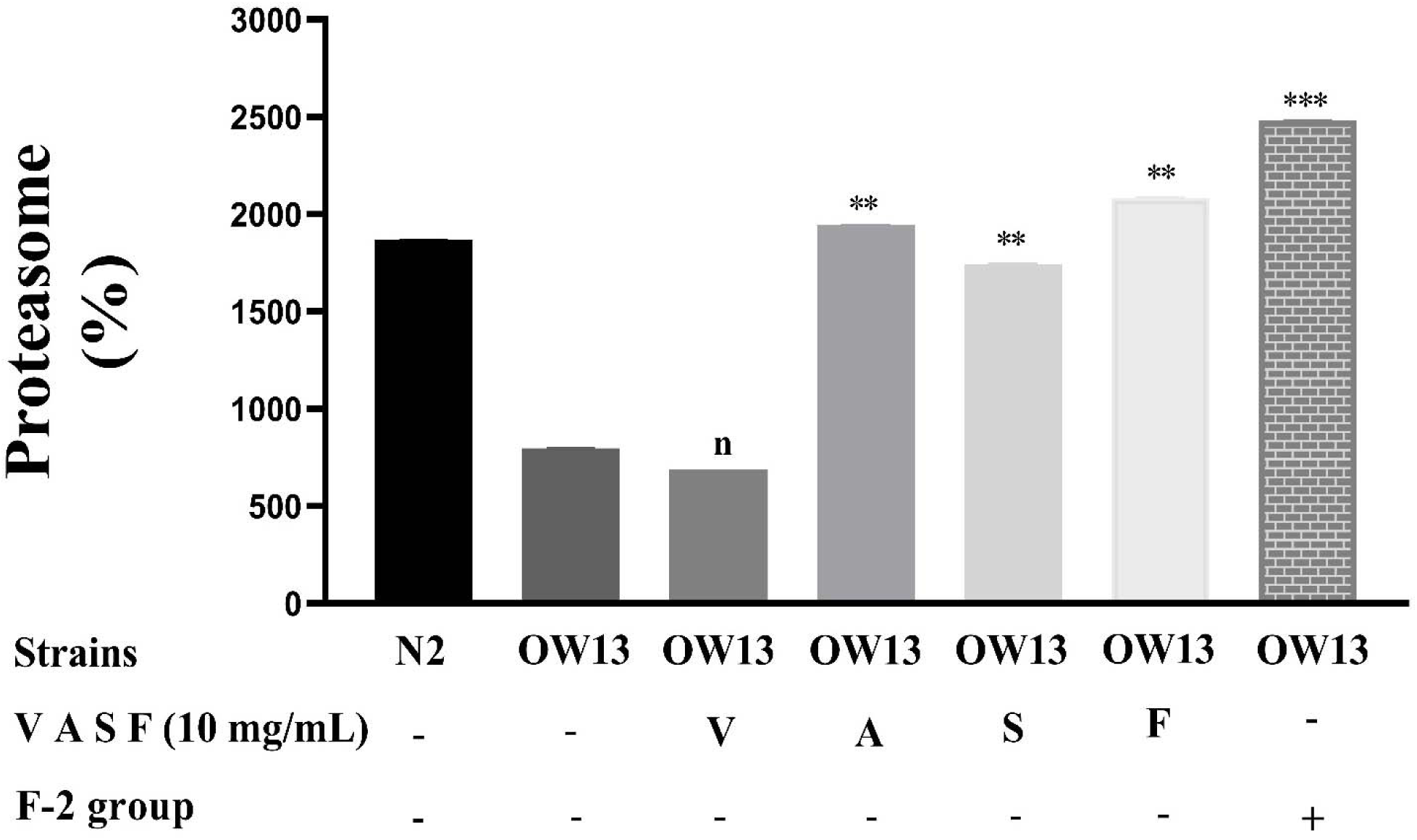
Formula drug F-2 exhibited better augmentation of ubiquitin-like proteasome activities than drug groups A, S, F on treatment in OW13. Proteasome activity was examined after 72 hrs of worm’s treatment with or without V, A, S, F at 10 mg/mL dilution, and F-2. Results showed that drug groups A, S, F on treatment significantly augmented the ubiquitin-like proteasome functions compared to control groups than drug V. Drug V is the only group that did not enhance the proteasome level in OW13 worms. The data was calculated by mean ± SD (n = 4). ns depicting no significant difference compared with control and V drug dilutions. ** shows the significant difference between A (62%, *p*≤0.005), S (55%, *p*≤0.005), F (68%, *p*≤0.005) treated and untreated groups at 10 mg/mL. At the same time, the F-2 (75%, *p*≤0.005) was treated in the similar way. F-2 significantly enhanced the ubiquitin-like proteasome activity much better than single drug treatment (A, S, F) than the control group’s *p*≤0.005, represented by ***. Together, these four drugs in F-2 showed more decisive neuroprotective actions than a single drug on treatment.

### 3.9 F-2 enhanced the ubiquitin-like proteasome activity higher than V, A, S, F in OW13 *C. elegans*

Several factors contributing to PD’s etiologic involve neuroinflammation, cellular apoptosis, autophagy insufficiency, and proteasome dysregulations ^35^. The ubiquitin-proteasome system is an essential feature of the proteostasis complex to degrade the internal misfolded cellular proteins ^36^. Consequently, to confirm whether ubiquitin-like proteasome enhanced on α-Syn reductions, we observed the V, A, S, F, and F-2 properties in OW13 worms and conducted a proteasome assay with the fluorogenic substrate (LLVY-R110-AMC). Our assessment showed that drug groups A (62%, *p*≤0.005), S (55%, *p*≤0.005), F (68%, *p*≤0.005) at 10 mg/mL, and F-2 (75%, *p*≤0.005) augmented the proteasome’s ubiquitin-like activity in the OW13 worms compared to untreated groups, except the drug V. F-2 exhibited 24.67 % better proteasome activity compared with the average of A, S, F drug groups. While the levels of proteasome activities in OW13 worms are about 56 % lesser than N2 worms. N2 is used here as a negative control group. These results proved that drug groups A, S, F and F-2 augmented the proteasome activity on treatment, but the formula drug F-2 exhibited higher proteasomes activity than single drug treatment (**Figure 7)**.

**Figure 7:**
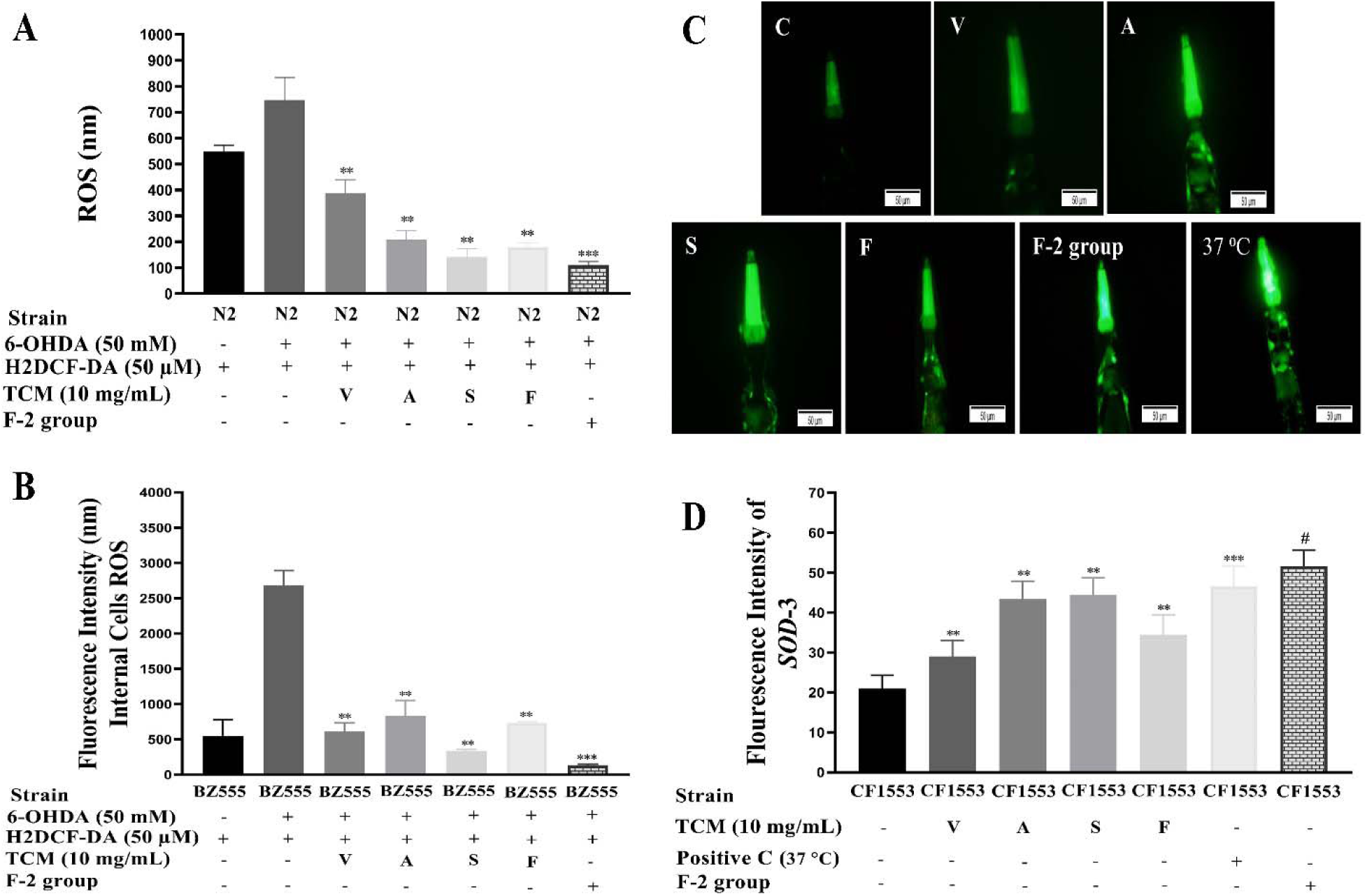
**(A&B)** Describing the antioxidative role of V, A, S, F and F-2. 6-OHDA exposed N2 and BZ555 *C. elegans* were treated with V, A, S, F at 10 mg/mL and F-2 for 72 hrs. Later, worms were treated with fluorogenic substrate H_2_DCF-DA (50 µM) in 96 wells black microtiter plates, and the reading was recorded. **(A)** Results showed that V, A, S, F significantly reduced the ROS productions in N2 worms compared to control groups. Similarly, **(B)** represents the internal cells’ antioxidative activities of V, A, S, F, and F-2 in BZ555 worm’s lysate on treatment. Data representing the mean±SD (n=10). ** showing the significant difference of (*p*≤0.005) between 6-OHDA treated group (control) vs V (43.3%, *p*≤0.005), A (60.7%, *p*≤0.005), S (68.2%, *p*≤0.005), F (64.5%, *p*≤0.005)/6-OHDA groups in both the worm’s (N2). Similarly for BZ555 on V (74%, *p*≤0.005), on A (68%, *p*≤0.005), on S (74%, *p*≤0.005), and on F (70%, *p*≤0.005). F-2 was treated the same way and calculated by mean±SD (n=10). F-2 reduced the ROS generation up to 78.34 % in N2 worms and 83.1 % in BZ555 compared to the control groups, much better than the V, A, S, and F better than single drug-treated groups (V, A, S, F)/6-OHDA and the control groups, *p*≤0.005. While **(C&D)** Reporter gene *SOD*-3::gfp increased the antioxidant enzymes in the CF1553 on V, A, S, F treatment. **(C)** Clearly showing the GFP over expressions of *SOD*-3 in the worms treated with V, A, S, F at 10 mg/mL compared to untreated groups. **(D)** Presenting the graphical form of *SOD*-3 reporter gene expressions results, showing that V, A, S, F enhanced the antioxidative enzyme actions while alleviating oxidative stress, a significant cause of PD. Data were calculated by mean+SD. Heat stress at 37 C to the worms was used as the positive control. ** describing the significant difference of *p*≤0.005 between V (18%, *p*≤0.005), A (38%, *p*≤0.005), S (39%, *p*≤0.005), F (23%, *p*≤0.005) treated and untreated groups. *** representing the significant difference between F-2 (45%, *p*≤0.005) and the control group. # representing the significant difference of *p*≤0.005 between the positive control and untreated groups.

### 3.10 F-2 reduced ROS better than V, A, S, F in DA neuron degenerative animals

Previously, the study proved that mitochondrial ROS plays a precarious part in degenerating DA neurons and escalations of α-Syn aggregative toxicity in PD ^37^. The aim was to detect the active antioxidative role of V, A. S. F and F-2. Our experimental assessment confirmed that V, A, S, F drug groups could decrease ROS levels in 6-OHDA exposed N2 and BZ555 *C. elegans* via antioxidative activities. In 6-OHDA-exposed worms, ROS significantly increased by about 24.76% in N2 and 68.43% in BZ555 compared to unexposed worms (*p*≤0.005). Hence, V (43.3%, *p*≤0.005), A (60.7%, *p*≤0.005), S (68.2%, *p*≤0.005), F (64.5%, *p*≤0.005) at 10 mg/mL significantly abridged the ROS generations in 6-OHDA exposed N2 worms. (**Figure 8A)** describes the results measured abruptly after H_2_DCF-DA transferring to the experimental groups, and reading was taken every 15 min for an hr. While (**Figure 8B)** represents the results counted after 1 h of incubation of transgenic BZ555 *C. elegans* lysate with H2DCF-DA, and reading was recorded every 10 min for 100 min. In 6-OHDA treated BZ555 worms, V drug reduces ROS up to (74%, *p*≤0.005), in A (68%, *p*≤0.005), in S (74%, *p*≤0.005), and in F (70%, *p*≤0.005). Our results verified that in both the worms V, A, S, F declined ROS generations in vivo. Similarly, the F-2 reduced the ROS productions in both the worms (up to 78.34 % in N2 worms and 83.1 % in BZ555 compared to the control groups, much better than the V, A, S, and F.

**Figure 8:**
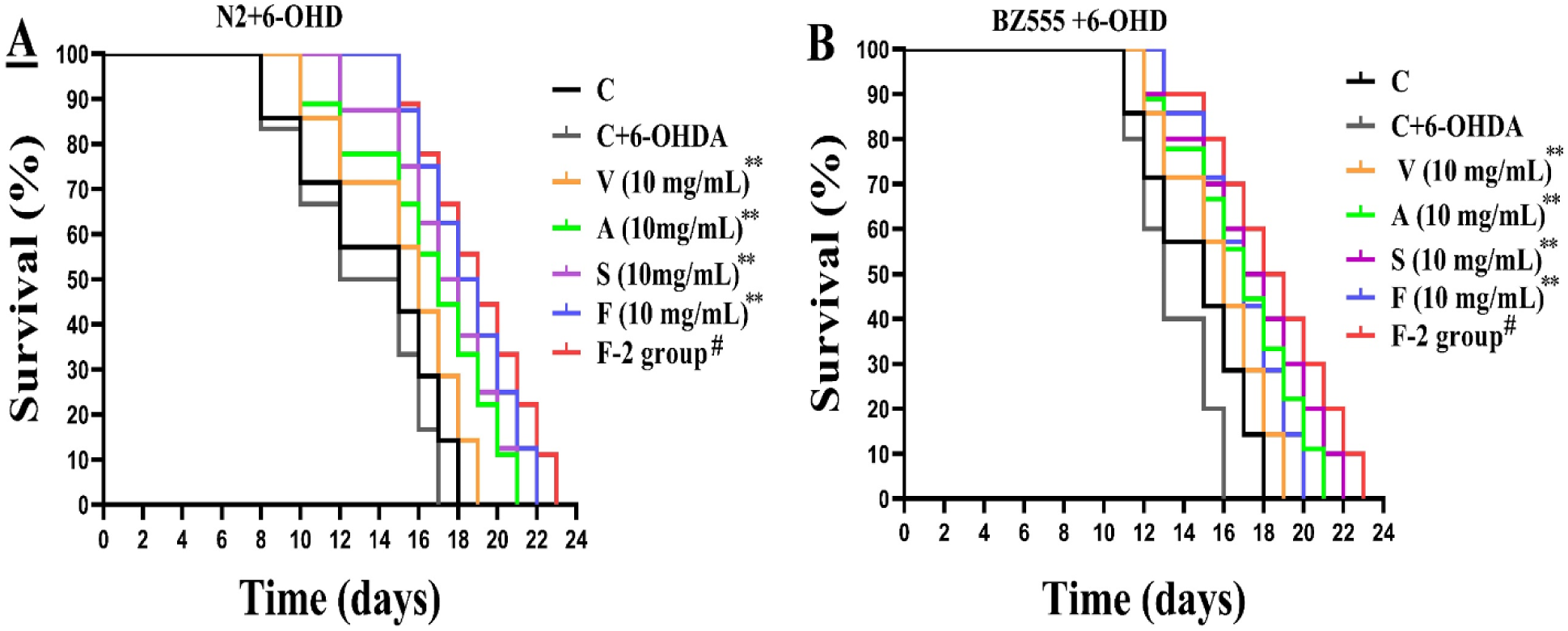
**(A&B)**. Representing the lifespan analysis results from the 6-OHDA exposed N2 and BZ555 worms treated with V, A, S, F at 10 mg/mL. Control was treated without drugs. Each drug remarkably expanded the nematodes’ lifespan up to 1.81 ± 1.21 d (V), 3.08 ± 0.45 d (A), 4.36 ± 0.71 d (S), 4.76 ± 1.33 d (F) in N2 and 2.03 ± 0.91 d (V), 3.01 ± 1.45 d (A), 3.79 ± 1.51 d (S), 4.83 ± 2.09 d (F) compared to the 6-OHDA treated groups. At the same time, the F-2 wa treated in the similar way and improved the worm’s lifespan up to 5.697 d in N2, and 6.89 d in BZ555,, (much higher than the single drug on treatment) when compared with the 6-OHDA treated (control) groups. Data was computed by Keplan Meir Survival test (n=3). ** describing the significant difference of *p*≤0.005 between V, A, S, F treated and untreated groups. # representing the significant difference of *p*≤0.005 between the F-2 and 6-OHDA treated groups. Significant diference were analyzed by survival surve comoarsion method in Graphpad.

### 3.11 F-2 enhanced the antioxidant (*SOD*-3) expressions greater than V, A, S, F in CF1553

Antioxidant enzymes like sodium superoxide dismutase (*SOD*-3) play an important part in decreasing ROS generations to enhance an organism’s lifespan ^38^. As it is known, PD is the second-most age-related neurological disorder. Mitochondrial ROS generations by oxidative stress is a significant risk factor that initiates PD. Excessive ROS generations have a prominent role in degenerating DA neurons and raising the α-Syn aggregations in the SNPc of the brain. Hence, we have evaluated the active role of the V, A, S, F and F-2 in augmenting the *SOD*-3 activities in CF1553 transgenic worms. *SOD*-3 managed oxidative stress via decreasing ROS and improved lifespans. On treatment, V (18%, *p*≤0.005), A (38%, *p*≤0.005), S (39%, *p*≤0.005), F (23%, *p*≤0.005) demonstrated a significant enhancement in *SOD*-3::GFP expression levels compared to control groups **Figure 8C**. Similarly, we treated the CF1553 worms with the F-2, and the results showed that F-2 enhanced the *SOD*-3 expressions by about (45%, *p*≤0.005) compared with control groups, much better than single drugs and positive control group (39.6%, *p*≤0.005). Our results specified that V, A, S, F, and F-2 drugs enhance longevity and reduce internal ROS generations when treated on *C. elegans*. *C. elegans* treated at 37 □C were considered a positive control (**Figure 8C&8D)**.

### 3.12 F-2 enhanced the lifespan of *C. elegans* longer than V, A, S, F

Aging is the main factor in developing neurodegenerative disorders and shortening life expectancy ^39^. Therefore, we investigated the effect of V, A, S, F and F-2 on the lifespans of 6-OHDA-exposed N2 and BZ555 *C. elegans*. The objective was to observe the differences between V, A, S, F, and F-2 effects on worm aging. 6-OHDA-treated N2 and BZ555 *C. elegans* showed smaller lifespans (16.08 ± 0.45 d) when compared to untreated *C. elegans* (18.01 ± 0.49 d,*p*≤0.005) via Kaplan Meier ^40^ survival cure analysis, signifying the toxic effect of 6-OHDA to hasten senescence. Therefore, V, A, S, F drug groups at 10 mg/mL prolonged the lifespan significantly (**Figure 9A&9B)**, delayed the 6-OHDA-induced senescence, and enhanced the worms’ lifespan up to 4.67 days. While, the F-2 was treated the similar way and promoted the life expectancies of worms up to 6.41 days compared to the 6-OHDA/Cont groups. Therefore, our experimental examinations verified that the F-2 stopped the aging in treated worms and enhanced the worm’s life durations up to 2.11 days longer than the average single-drug treatment, approximately similar in both worms **(Table 4)**. Table 4 explains the average lifespan of worms to the percentage of increase lifespan after V, A, S, F, and F-2 treatment. Worm’s average lifespan was deduced after taking the average of three biological repeats of each drug dilution of experiments and later evaluate by KM survival analysis. While % of increased lifespan has been calculated by a mathematical formula given below.

*% worms lifespan increase = (6-OHDA (V, A, S, F and F-2) ÷ 6-OHDA treated worms × 100) 100*

**Table 4:**
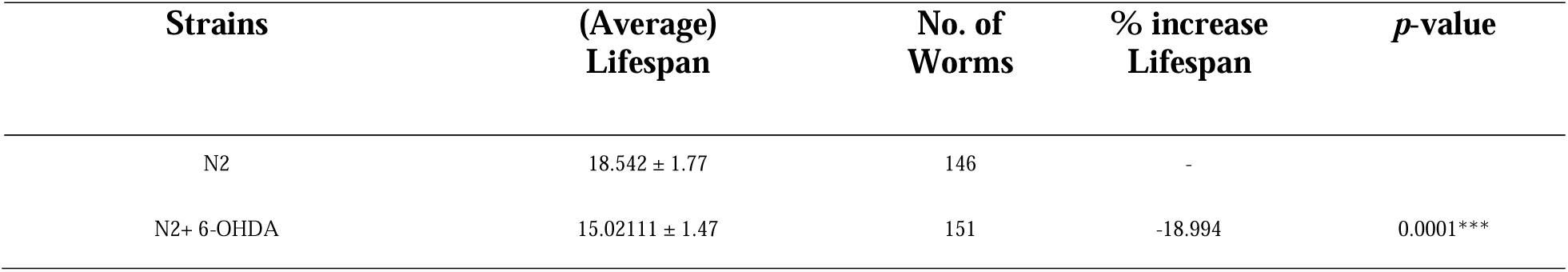
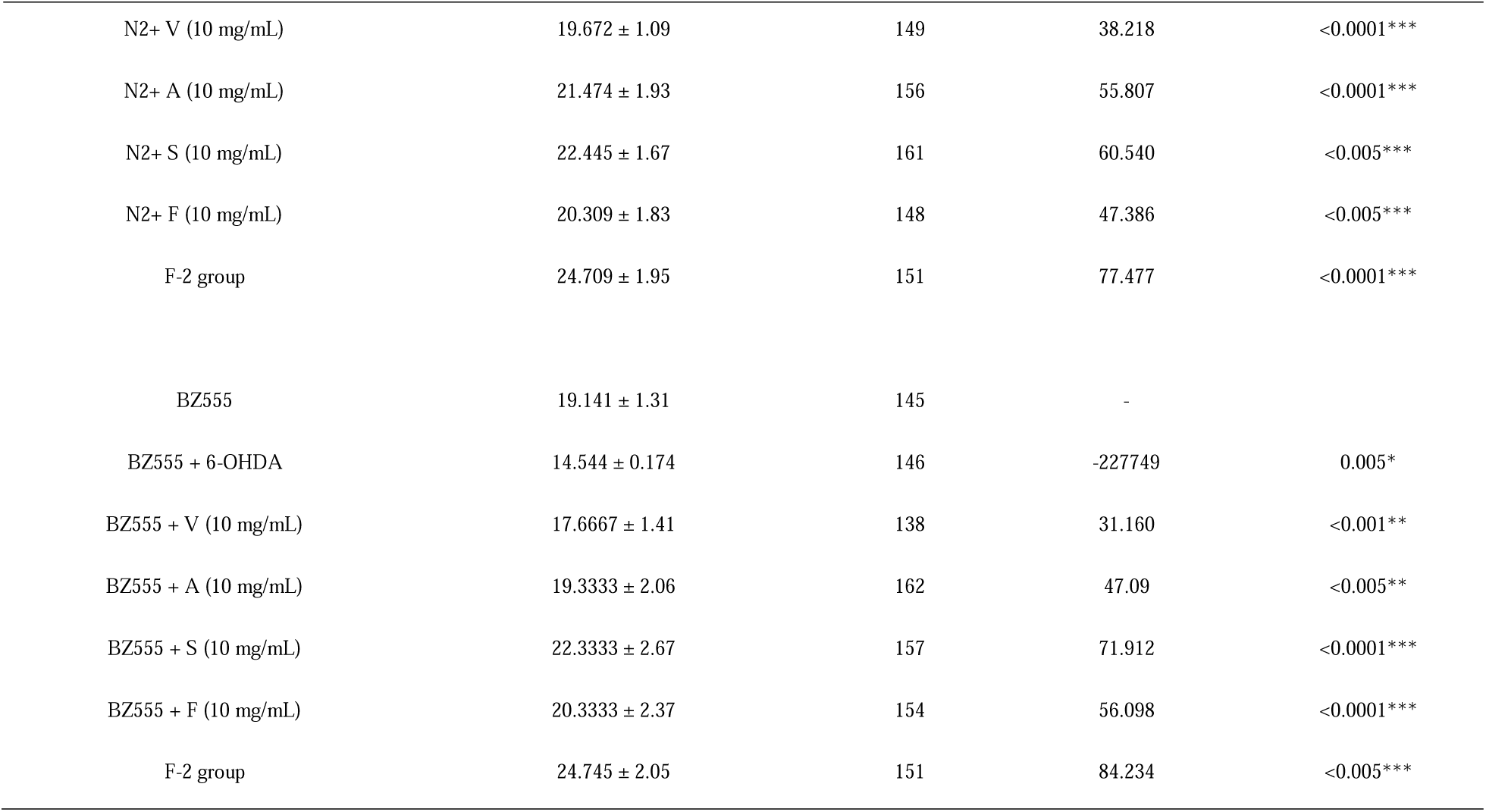
Lists of strains, mean lifespan, numbers of worms, percentage of increased lifespan, and *P*-values of N2, BZ555 after treatments with V, A, S, F at 10 mg/mL and F-2.

## 4 Discussion

Zhi-Shi-Huang-Wu is a formula computed in current study from four best HSP 70 promoter activator TCMs, which comprises Valeriana jatamansi (V, with Chinese herbal name “zhi zhu xiang”), Acorius talarinowii (A, with Chinese herbal name “shi chang pu”), Scutellaria baicalensis (S, with Chinese herbal name “huang qin”), and Fructus Schisandrae (F, with Chinese herbal name “wu wei zi”) at 8:4:2:1 ratio, called F-2, calculated after orthogonal experimentations of **Table 3** and has multiples therapeutic actions. Orthogonal experimentations were conducted in transgenic OW13 and CF1553 integrated with protein α-Syn tagged with GFP promoter and antioxidant *SOD*-3 genes tagged with GFP expressed in muscles and head regions, respectively to computed F-2 formula drug (results are given in supplementary data).

TCM has a long history of using herbal medicine in central Asia, especially in China, Kora, Japan, and India, to cure multiple human diseases. Traditionally, each TCM has shown biological activities to cure human disorders, like liver damage, respiratory inflammations, toxic effects, sedatives, epilepsy, and nervous system problems. We have selected four TCM out of 35 on their screening-based study against heat shock protein (HSP) promoter activations in the present study **Table 1**. Mentioned four drugs significantly enhanced the HSP 70 promoters in animals on treatment in our previous study [Wang Yajun. Anti-AD traditional Chinese medicine screening based on the promoter of heat shock protein HSP70, Lanzhou University, 2017]. HSP can prevent false folding and protein aggregation, especially the HSP 70 ^41^. While, HSP 70 plays a vital role in reducing the accumulation of α-Syn, Aβ, oxidative stress, and inhibiting nerve inflammations ^42, 43^. Among the four TCM, V, and S is the best HSP 70 promoter activator which can activate the HSP 70 expressions in pGL3-HSP70 and pRL-TK co-transfected HEK-293T cells as low as 0.4 mg/mL decoction, while others are 3.2 mg/mL (A), 1.2 mg/mL (S), 5.5 mg/mL (F), respectively. Therefore, we included the V drug in the four best HSP 70 promoter activator TCM and treated them as anti-PD drugs. Unluckly, V failed to prove be an anti-PD drug on treatment. Because of V drug HSP 70 promoter activation and antioxidant properties, we have included V with A, S, F in F-2. Later, we have developed experimental strategies to evaluate the neuroprotective role of the F-2. The V, A, S, F extracts have been reported with major constituents comprised of flavonoids, triterpenoids, phenols, and glycosides. This study has selected transgenic *C. elegans* as PD models. These *C. elegans* based analyses could be effective for rapid assessments, cheaper, and scrutinizing new neuroprotective drugs in a short time ^40^. OP50 (*E. coli*) strains are used as a food source for *C. elegans* models. According to previous research, few drugs (extracts and their components) have anti-bacterial properties and showed minimum inhibitory concentration (MIC) against bacterial strains ^44^. To observe the MIC and toxic effects of TCM extracts. We performed a food clearance test of V, A, S, F and F-2 to measure the non-toxic concentrations, in transgenic BZ555, OW13, and wild-type N2 worms. N2 was considered a negative control group to differentiate the V, A, S, F and F-2 cytotoxicity behavior from transgenic *C. elegans*. Result confirmed that S, F drug groups exhibited toxicity at 20 mg/mL and 30 mg/mL to the treated *C. elegans,* caused non-motility and even death within time periode. Therefore, we have considered the non-toxic concentrations for conducting experiments in *C. elegans* up to 10 mg/mL.

DA neuron degenerations are the leading cause of PD. In this research, we used 6-OHDA as a neurotoxic to induce degenerations in DA neurons of BZ555. BZ555 worms with Pdat-1 promoter tagged by DA-GFP ^45^. In vivo, 6-OHDA uptake causes a desensitization impact on DA neuron level to generate PD-like symptoms in BZ555 ^46^. After generating the self-neurodegenerative models, we treated them with V, A, S, F, and F-2. A, S, Fsignificantly recovered the 6-OHDA generated DA neuron degenerations, except drug V. Similarly, F-2 formula drug can diminish DA neuron degenerations much better than A, S, F. The mediated DA neuronal circuits control the basal slowing rate of *C. elegans* via sensing the food ^47^. A degenerative DA neuron decreases the dopamine level, leading the nematodes to be imperfect in perceiving foods. Therefore, we observed the food sensing behavior test in both the worms N2 and BZ555, intoxicated with 6-OHDA. The objective was to observe whether V, A, F, S and F-2 could save the food-sensing behaviour of nematodes under 6-OHDA treatment or not. On treatment, we observed bending frequency of N2 mainly decreases to 61% in the bacterial lawn compared to 6-OHDA treated N2 nematodes, while 63.71% in BZ555. N2 was used as a negative control group to monitor the behavioural differences in transgenic and wild-type worms. Except for drug group V, our results verified A, F, S, and F-2 restored food sensing behavior in N2 and BZ555 *C. elegans* after 6-OHDA treatment, dose-dependently.

6-OHDA intoxications cause generations of free radicals ^48^. These free radicals are the major contributor to the generations of mitochondrial ROS ^49^. Previous studies proved that mitochondrial ROS played a vital role in degenerations of DA neurons and Lewy bodies’ formations. Lewy bodies are the aggregative product of α-Syn, which is the main culprit to cause PD ^50^. While aggregated α-Syn in the muscles walls identifying the in vivo toxicity leads to failure of PD-like progressive motility in OW13 ^51^. Perhaps, α-Syn activities mitigated the PD dysregulations and later downregulated the self-machinery destructions. The precise mechanism for these results will need advanced investigations. Moreover, A, S, F, and F-2 formula drug abridged the α-Syn aggregative toxicity in OW13 transgenic worms’ dose-dependently, except the group V, verified by western blotting analysis.

Large quantities of lipids entities in the CNS propose that their existence is not restricted to the cell’s drive and structural components formations. Some lipids molecules in the CNS are acquainted with play a vital part in neurotransmission. Hence, all cellular signalings activity controlled by lipids molecules can change the protein’s scaffoldings and locations via microdomain membrane control centres ^52^. This protective role of the A, S, F, and F-2 drug groups reduces lipids peroxidation and protects lipid disturbed functionality to keep a well-organized cellular signaling pathway. A, S, F and F-2 formula drug enhanced lipid depositions in OW13 worms, assessed after staining with Nile Red dye. Our outcomes confirmed that the α-Syn anti-accumulated effect of A, S, F and F-2 might activate antiapoptotic machinery to fight the aggregated misfolded proteins via activating the HSP 70 promoter and mediating the ubiquitination proteasome systems (UPS). The UPS endorses cell survival on damage under environmental stress conditions ^53^. Besides, our ubiquitin-like proteasome activity assay demonstrated that A, S, F, and F-2 drug groups actively enhanced the proteasome 20S functions on treatment in transgenic worms. Excessive ROS production induces oxidative stress, which is one of the leading causes of PD. Therefore, we observed the antioxidative properties of V, A, S, F, and F-2 in N2 and BZ555. Our results confirmed that V, A, S, F, and F-2 formula drug extracts reduced ROS generations at live and internal cells level in *C. elegans* on treatment. Previously it was shown that antioxidants superoxide dismutases (*SOD*-3) expressions stopped the free radicals during the cell’s oxidative reactions and stopped the aging process. Hence, drugs V, A, S, F, and F-2 promoted the *SOD*-3 expressions in transgenic CF1553 to stop the ROS injuries and reduce oxidative stress. As we know, PD is an age-related disorder ^54^. Therefore, we have done a life expectancy assay to examine the effects of the V, A, S, F, and F-2 on worm lifespans. Our results confirmed that V, A, S, F increase the age expectancy by about 4 days in 6-OHDA intoxicated worms compared with control groups. Similarly, the F-2 formula drug prolonged the worm’s lifespans 2.11 days longer than individual drugs V, A, S, F groups.

As per our knowledge, it is the first study to report neuroprotective properties of V, A, S, F drug extract against PD in transgenic *C. elegans* models. The purpose was to observe how these 4 selected drugs (V, A, S, F) in specific proportions (F-2) protects DA neuronal damage, reduced α-Syn aggregative toxicity, and stopped PD-like symptoms in treated worms. Our results showed that the neuroprotective effects of the F-2 are better than single drug groups on treatment. Therefore, we can say that these four drugs significantly supported and enhanced their neuroprotective functions against PD transgenic models on treatment. The neuroprotective mechanisms of action of V, A, S, F drugs extract and F-2 might follow the oxidative or ubiquitinated pathway, but further research is needed to be done in the future.

## 5 Conclusion

In conclusion our current study demonstrated that, Zhi-Shi-Huang-Wu in short F-2 formula improved the pathaology of PD much better than V, A, S, F resulting from curtaining the α-Syn accumulations, recovered 6-OHDA exposure on DA neurons, augmented the lipid deposits, proteasomes, and antioxidants (*SOD*-3) activities in transgenic *C. elegans*. Morever F-2 decreased the ROS generations in vitro and in vivo and prolonged the lifespan. Therefore, these finding strongly supported that F-2 may have considerable neuroprotective applications for PD via activations of the antioxidative pathway. In additions, further research will be required on F-2 to validate the anti-PD effects and explore the molecular mechanisms of action clinically in our future projects.

## List of abbreviations

PD: Parkinson’s disease
TCM: Traditional Chinese Medicine
α-Syn: Alpha-synuclein
6-OHDA: 6-hydroxydopamine
CGC: Caenorhabditis Genetics Centre
NGM: Nematode growth medium
OD: Optical density
ROS: Reactive oxygen species
DA Neuron: Dopaminergic Neuron
SNPc: substantia nigra pars compacta
SOD: Superoxide dismutase
UPS: Ubiquitin-like Proteasome System

## Acknowledgements

We sincerely thank all the participants who took part in this study; in particular, Dr. Liu Yan gratefully acknowledged her assistance in writing and guidance in experimental work. This exploratory study was funded by the Natural Science Foundation of Gansu Province of China, 20JR10RA596, 20JR10RA756, and the Talent innovation and entrepreneurship project of Lanzhou City 2020-RC-43.

## Declarations Ethical statement

n/a

## Consent for publication

n/a

## Authors’ contributions

Fahim M conducted the experimental work, Yan Liu guided the practical work and helped in writing, Ningbo Wang encouraged experimentations, Longhe Zhao supported in conducting the research, Yongtao Zhou and Hui Yang financially supported in achieving this study project, and Hongyu Li supervised the project. All authors contributed to the concept of this study. All authors provided critical feedback and helped shape the research, analysis, and manuscript.

## Availability of data and materials

The aggregate data supporting findings contained within this manuscript will be shared upon request submitted to the corresponding author.

