## Supplementary material for "Zhi-Shi-Huang-Wu slows Parkinson’s progression in transgenic *C. elegans* models": Supplemetary data

**Supplementary experimental data**

### 1 HPLC analysis of Traditional Chinese Medicine (V, A, S, F)

#### 1.1 HPLC required Reagents.

Acetonitrile 80 % (HPLC grade) and Ultrapure water (20 %) were obtained using the Milli-Q system (Millipore, Bedford, MA, USA) was used in the experiments. Methanol (HPLC grade) was used to clean the column. Eight standards for quantitative analysis were purchased from the National Institution for Food and Drug Control.

### 2 Results

#### 2.1 Wavelength optimization.

The eight mentioned standards were scanned using the UV detector between 190 and 400 nm. The maximum absorbance values of Caffic acid, Valtrate, β-asarone, α-asarone, Gallic acid, Schizandrol A, Baicalin, and Chrysin were 314, 310, 270, 280, 330 nm, 254, 280, and 370, respectively. Given the serious end absorption near 270 nm, the detection wavelength was set at 270 nm, in which all of the compounds have an apparent absorption band. The results are shown in supplementary figure 1**.**

#### 2.2 Detector optimization.

The UV detector can detect only one wavelength at a time. Therefore, measuring the standard's characteristic absorption and retention time was easier. A general assumption can be made using the characteristic absorption and retention time. In this way, the error rate is unusually reduced and suitably used to valuation each chromatographic peak of the corresponding samples.

#### 2.3 Mobile phase optimization.

Several organic solvents in different ratios resulted in different peak separations. The composition and ratios of the mobile phase were optimized using different organic solvents, including acetonitrile and water in different concentrations. Methanol was used to clean the column. In the present study, the mobile phase composition was selected as the mobile phase for the simultaneous analysis because of better resolution and shorter analysis time than other phases. Typical HPLC-PDA chromatograms of the standard samples are presented in (Supplementary figure 1).

#### 2.4 Stability.

The mixed working standard solutions of different concentrations were analyzed at 0, 3, 6, 9, 15, 21, and 24 h at room temperature through the method's stability. The peak area was recorded, and RSD was calculated. The RSDs were 0.30%, 0.31%, 0.30%, 0.37%, and 0.48%, 0.54 %, 0.67 % and 0.71 % (n= 8), which indicated that the established method was stable in a day.

#### 2.5 Repeatability.

Two portions were precisely weighed from each V, A, S, F and formula drug sample; the Caffic acid, Valtrate, β-asarone, α-asarone, Gallic acid, Schizandrol A, Baicalin, and Chrysin contents were detected through the developed method; the RSDs of the samples were 2.80 %, 7.284 %, 3.39 %, 3.29 %, 2.8 4%, 3.19 %, 2.81 % and 3.22 % respectively. Standard contents were .246 mg/mL, .280 mg/mL, .270 mg/mL, .253 mg/mL, .293 mg/mL, .262 mg/mL, and 0.26 mg/.253 mg/mL, respectively.

#### 2.6 Applications.

Using the new and modern method, the analytic contents of the four selected TCM and formula drugs from various regions of China were measured. Marked differences were observed in the analytic content of TCM samples. This TCM has the highest amount of Valtrate (7.284 %) in V, α-asarone (3.297 %) in A, Schizandrol A (3.19 %) in F, and Baicalin (2.88 %) in S were relatively high. The analytic entities were almost similar to the other area of herbs (Supplementary figure 1).

#### 2.7 Results

##### 2.7.1 Wavelength Optimization.

The eight mentioned standards were scanned using the UV detector between 190 and 400 nm. The maximum absorbance values of Caffic acid, Valtrate, β-asarone, α-asarone, Gallic acid, Schizandrol A, Baicalin, and Chrysin were 314, 310, 270, 280, 330 nm, 254, 280, and 370, respectively. Given the serious end absorption near 270 nm, the detection wavelength was set at 270 nm, in which all of the compounds have an apparent absorption band. The results are shown in Figure 1**.**

##### 2.7.2 Detector Optimization.

The UV detector can detect only one wavelength at a time. Therefore, measuring the standard's characteristic absorption and retention time was easier. A general assumption can be made using the characteristic absorption and retention time. In this way, the error rate is unusually reduced and suitably used to valuation each chromatographic peak of the corresponding samples.

##### 2.7.3 Mobile Phase Optimization.

Several organic solvents in different ratios resulted in different peak separations. The composition and ratios of the mobile phase were optimized using different organic solvents, including acetonitrile and water in different concentrations. Methanol was used to clean the column. In the present study, the mobile phase composition was selected as the mobile phase for the simultaneous analysis because of better resolution and shorter analysis time than other phases. Typical HPLC-PDA chromatograms of the standard samples are presented in (supplementary figure 1).

##### 2.7.4 Stability.

The mixed working standard solutions were analyzed at 0, 3, 6, 9, 15, 21, and 24 h at room temperature to evaluate method stability. The peak areas of eight analytes were recorded and the relative standard deviations (RSDs) were calculated. The RSD values ranged from **0.30% to 0.71% (n = 8)**, indicating that the developed method exhibited good stability within 24 h.

| Compound | RSD (%) (n=8) |
| --- | --- |
| Caffeic acid | 0.30 |
| Valtrate | 0.31 |
| β-Asarone | 0.30 |
| α-Asarone | 0.37 |
| Gallic acid | 0.48 |
| Schizandrol A | 0.54 |
| Baicalin | 0.67 |
| Chrysin | 0.71 |

##### 2.7.5 Repeatability.

Two portions of each sample (V, A, S, F, and the formula drug) were accurately weighed and analyzed using the developed HPLC method. The contents of caffeic acid, valtrate, β-asarone, α-asarone, gallic acid, schizandrol A, baicalin, and chrysin were determined. The RSD values ranged from 2.80% to 7.28%, demonstrating acceptable repeatability of the method.

| Compound | Standard Concentration (mg/mL) | RSD (%) |
| --- | --- | --- |
| Caffeic acid | 0.246 | 2.80 |
| Valtrate | 0.280 | 7.28 |
| β-Asarone | 0.270 | 3.39 |
| α-Asarone | 0.253 | 3.29 |
| Gallic acid | 0.293 | 2.84 |
| Schizandrol A | 0.262 | 3.19 |
| Baicalin | 0.260 | 2.81 |
| Chrysin | 0.253 | 3.22 |

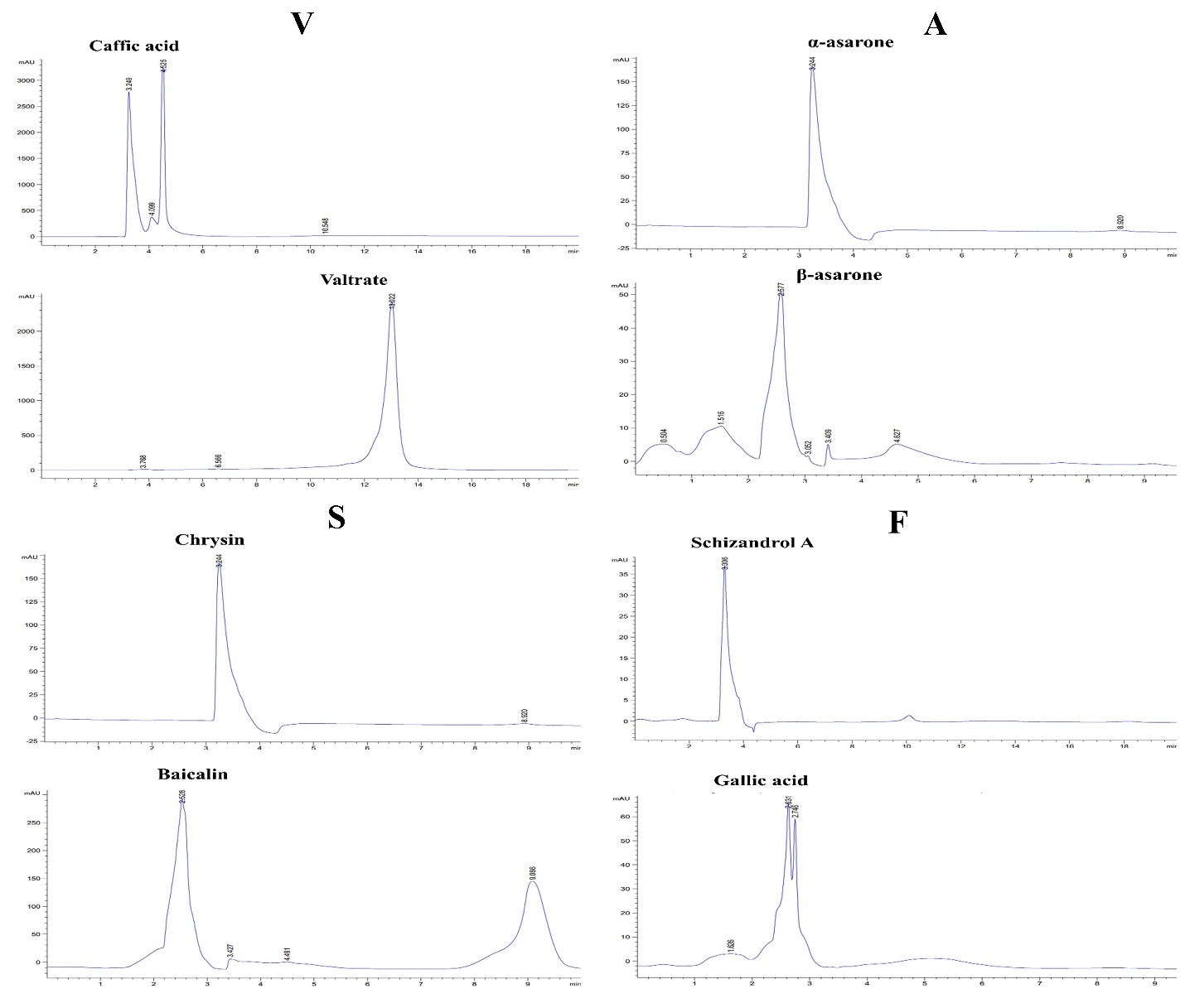
**Supplementary figure 1:** Explaining the chromatograms of particular standards for each TCM Caffic acid, Valtrate, β-asarone, α-asarone, Gallic acid, Schizandrol A, Baicalin, and Chrysin under optimal chromatographic conditions.

##### 2.7.6 Applications.

Using the new and modern method, the analytic contents of the four selected TCM and formula drugs from various regions of China were measured. Marked differences were observed in the analytic content of TCM samples. This TCM has the highest amount of Valtrate (7.284 %) in V, α-asarone (3.297 %) in A, Schizandrol A (3.19 %) in F, and Baicalin (2.88 %) in S were relatively high. The analytic entities were almost similar to the other area of herbs (Supplementary figure 2).

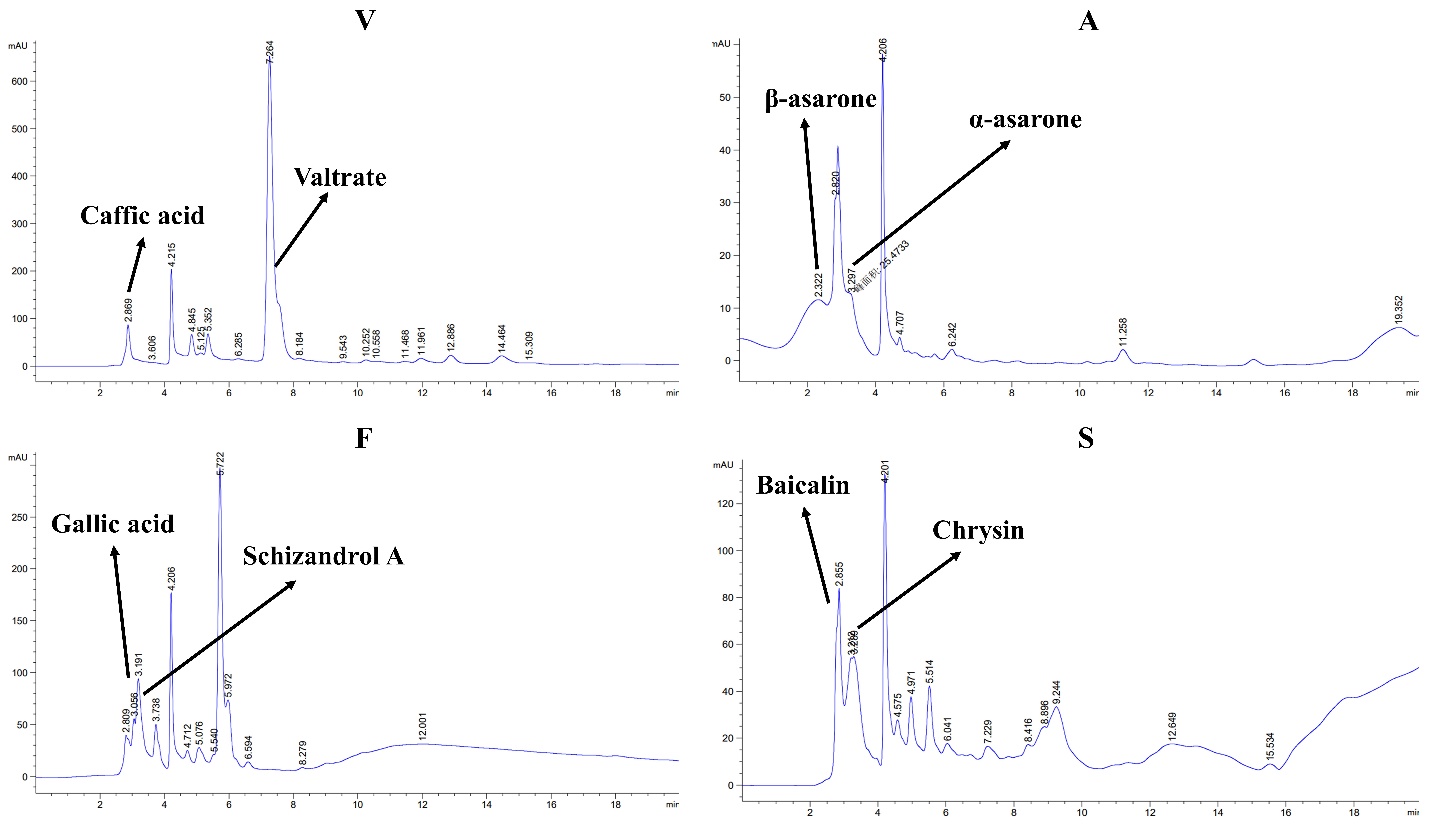
 **Supplementary figure 2:** Representing the chromatograms of V, A, F, S TCM with the percentage of standard-specific standards represented by peak values.

### 3 V, A, S, F Orthogonal experiments for α-Syn, lipid depositions and anti-oxidative expressions of *SOD-3* enzymes for calculating F-2 afficacy

Orthogonal experiments were performed according to the drug dilutions in **Table 3** to prepare a highly effective neuroprotective formula drug (F-2) to treat transgenic *C. elegans* models of PD. On our experimental results are given in (**supplementary figure 1AB-CD and 1E-1F**). We concluded that the drug V, A, S, F in Zhi-Shi-Huang-Wu formula at 8:4:2:1 ratio (specifically at V: 20 mg/mL, A: 10 mg/mL, S: 5 mg/mL, F: 2.5 mg/mL) named as formula-2 (F-2).

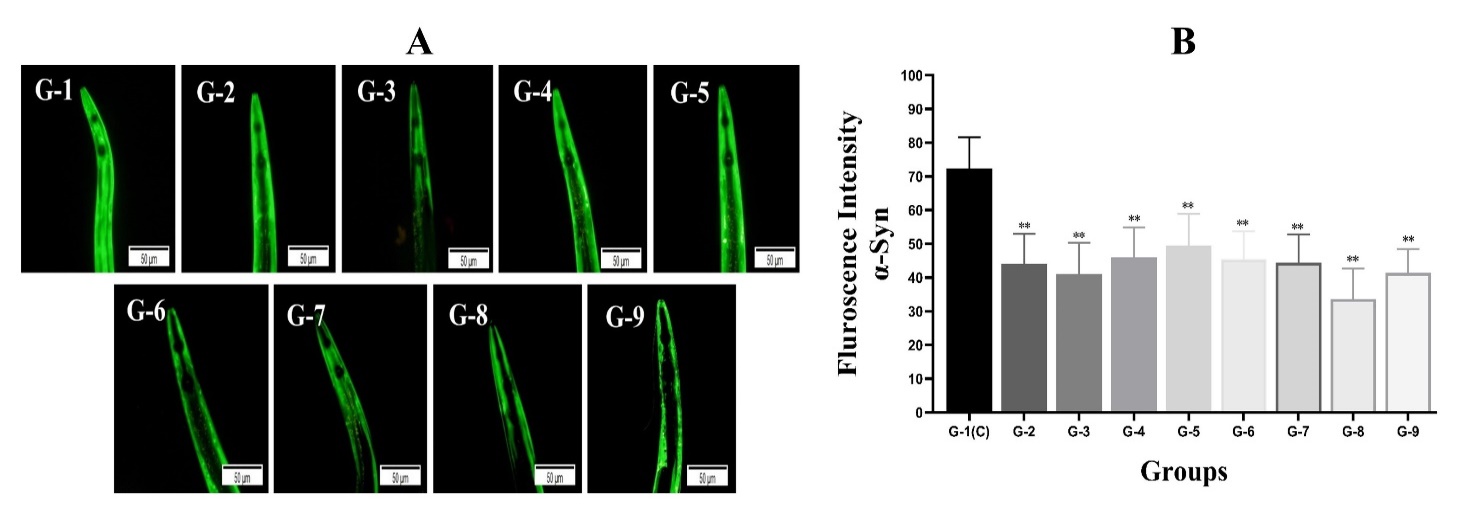

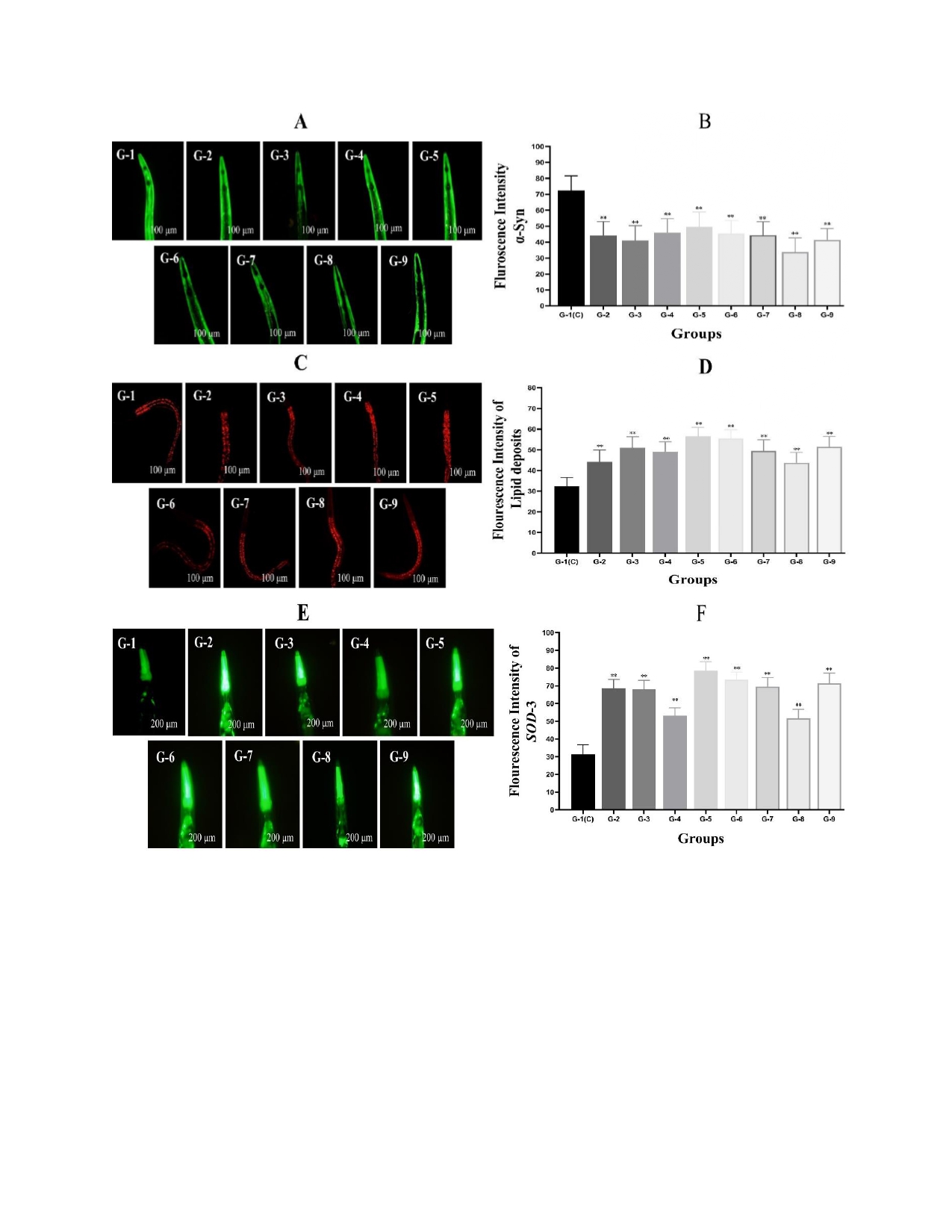

**Supplemenatry figure 3.** (A&B) Depicting the results of orthogonal drug dilutions in Table 2 from G-1-G-9 on treatment reduced the of α-Syn accumulation in OW13. (A) YFP fluorescence expressions in the muscles of worms OW13. The scale bar was 50 μm. (B) Bar Graph representing the quantitative data of the fluorescence intensities in OW13 after G-1🡪G-9 treatment. G-1 group treated without drug (control). ImageJ software was used to enumerate the fluorescence intensities of images. Data were computed by mean±SD (n = 3). More than 30 images were quantified in each experimental group. ** shows a significant difference between the drugs treated G-2 (34.3%), G-3 (40.6%), G-4 (36.1%), G-5 (29.4%), G-6 (33.5%), G-7 (36.2%), G-8 (67.3%), G-9 (55.4%) and untreated groups (G-1) (p≤0.005). Similarly we treated the OW13 with the following G-1-G-9 dilution for enhanced lipid deposits activity after stained with dye Nile Red **(3C)** Representative images are the Nile red staining of OW13 worms after G-1🡪G-9 treatment for 72 hrs. The scale bar was 100 μm. **(3D)** A graphical picture of fluorescence intensities of strain OW13 was stained with the Nile Red and quantified by software ImageJ. The data was calculated by mean ± SD (n = 30). G-1 group treated without drug. ImageJ software was used to enumerate the images fluorescence intensities. ** shows a significant difference between the drugs treated G-2 (23.4%), G-3 (29.6%), G-4 (28.4%), G-5 (32.3%), G-6 (31.8%), G-7 (30.8%), G-8 (27.1%), G-9 (31.4%) and untreated groups (G-1) (*p*≤0.005). Similarly, **(3E)** describing the superoxide dismutase enzymes expressions enhancement in CF1553 transgenic worms after G-1🡪G-9 treatment for 72 hrs. *SOD*-3 enzymes involved in reducing ROS and also enhanced the worm’s lifespan on expressions. The scale bar was 200 μm. **(3F)** A graphical picture of fluorescence intensities of strain CF1553 was quantified by software ImageJ. The data was calculated by mean ± SD (n = 30). G-1 group treated without drug. ImageJ software was used to enumerate the fluorescence intensities of images. ** shows a significant difference between the drugs treated G-2 (51.1%), G-3 (49.5%), G-4 (38.3%), G-5 (65.4%), G-6 (59.3%), G-7 (55.4%), G-8 (37.2%), G-9 (61.7%) and untreated groups (G-1) (*p*≤0.005).
